# Lymph node-targeted vaccine boosting of TCR-T cell therapy enhances anti-tumor function and eradicates solid tumors

**DOI:** 10.1101/2022.05.05.490779

**Authors:** Dylan J. Drakes, Abdulraouf M. Abbas, Jacqueline Shields, Martin P. Steinbuck, Aniela Jakubowski, Lochana M. Seenappa, Christopher M. Haqq, Peter C. DeMuth

## Abstract

While T cell receptor (TCR)-modified T cell therapies have shown promise against solid tumors, overall therapeutic benefits in clinical practice have been modest due in part to suboptimal T cell persistence and activation in vivo, alongside the possibility of tumor antigen escape. In this study, we demonstrate an approach to enhance the in vivo persistence and activation of TCR-T cells through combination with Amphiphile (AMP)-vaccination including cognate TCR-T peptides. AMP-modification improves lymph node targeting of conjugated tumor immunogens and adjuvants, thereby coordinating a robust T cell-priming endogenous immune response. Vaccine combination with TCR-T cell therapy provided simultaneous in vivo invigoration of adoptively transferred TCR-T cells and in situ priming of the endogenous anti-tumor T cell repertoire. The resulting induction of an adoptive and endogenous anti-tumor effect led to durable responses in established murine solid tumors refractory to TCR-T cell monotherapy. Protection against recurrence was associated with antigen spreading to additional tumor-associated antigens not targeted by vaccination. Enhanced anti-tumor efficacy was further correlated with pro-inflammatory lymph node transcriptional reprogramming and increased antigen presenting cell maturation, resulting in TCR-T cell expansion and functional enhancement in lymph nodes and solid tumor parenchyma without lymphodepletion. In vitro evaluation of AMP-peptides with matched human TCR-T cells targeting NY-ESO-1, mutant KRAS, and HPV16 E7 illustrated the clinical potential of AMP-vaccination to enhance human TCR-T cell proliferation, activation, and anti-tumor activity. Taken together, these studies provide rationale and evidence to support clinical evaluation of the combination of AMP-vaccination with TCR-T cell therapies to augment anti-tumor activity.

**Summary:** AMP-vaccination targets the lymph nodes to enhance TCR-T cell therapy resulting in solid tumor eradication and durable protection against recurrence.

## Introduction

Recent translational advancement of engineered T cells genetically modified to express tumor-specific TCRs has demonstrated the promise of this strategy to treat a variety of cancers (*1–9*). In contrast to chimeric antigen receptor (CAR) T cells, TCR-modified T cells (TCR-T cells) have been developed to target a variety of intracellular and extracellular tumor antigens in association with MHC suggesting the potential for wide-ranging therapeutic benefit resulting from tumor recognition with attractive fidelity and sensitivity (*8–15*). However, while recent clinical trial results have provided important proof-of-concept including demonstration of objective anti-tumor activity, the overall benefit to patients has been modest, particularly in solid tumor settings of therapy, where low response rates and poor durability remain significant challenges. Cell-intrinsic strategies to improve efficacy have included TCR affinity enhancement which may improve activity but often at the expense of dose-limiting adverse safety effects (*4, 8, 16*). Additional studies have suggested that instead of suboptimal TCR binding affinity, treatment failures may be attributed to a deficiency of in vivo T cell activation, poor initial engraftment, limited persistence, as well as multiple weaknesses in T cell anti-tumor functionality. Therefore strategies to promote the delivery of comprehensive stimulatory signals to TCR-T cells in vivo may be critical to enabling further clinical success (*10, 17*).

Numerous methods have been reported for enhancing T cell function in vivo, including direct genetic encoding of pro-inflammatory molecules promoting aspects of T cell-intrinsic or anti-tumor immune biology, provision of these agents given through combination therapy, or equipping T cells with exogenous sources of supportive or modulating factors via cell-modification with nanocarriers or other means (*18–21*). While these strategies have demonstrated potential to improve efficacy, they present a limited opportunity to supplement the inherently complex and multi-factored biology underlying the endogenous mechanisms contributing to effective T cell priming critical to effective anti-tumor responses. Concerted promotion of these mechanisms in vivo, including stimulation of the TCR-signaling complex, delivery of co-stimulation, and cytokine support, can be achieved through vaccination which potently engages the innate immune response to simultaneously deliver the comprehensive and complex suite of signals needed to activate and support the robust activation and function of effector T cells. Administration of TCR-specific epitope-based vaccines is highly attractive for this purpose, offering simple TCR-informed design of cognate peptide immunogens. However, as a result of their small molecular size, conventional peptides are known to be poorly immunogenic due to ineffective accumulation and retention in lymph nodes where adaptive immunity is orchestrated by resident antigen presenting cells (APCs) (*22, 23*). Previous studies have demonstrated the development of albumin “hitchhiking” utilizing Amphiphile (AMP)-conjugated vaccines that promote increased stability and specific trafficking and retention of vaccine payloads into lymph nodes (*23–26*). In this manner, both peptide antigens and pro-inflammatory molecular adjuvants can be coupled to lipophilic, albumin-binding domains, that non-covalently associate with endogenous tissue albumin at the injection site after administration (Fig. 1A). These AMP-albumin complexes restrict entry of vaccine components to the systemic blood circulation and resulting distribution away from lymphatics, preferentially re-targeting albumin-bound agents to the lymph nodes. Resulting uptake by resident APCs can subsequently initiate concerted presentation of cognate peptides and costimulatory ligands with cytokine secretion to potently activate and license T cells in vivo (*27*).

**Fig. 1:**
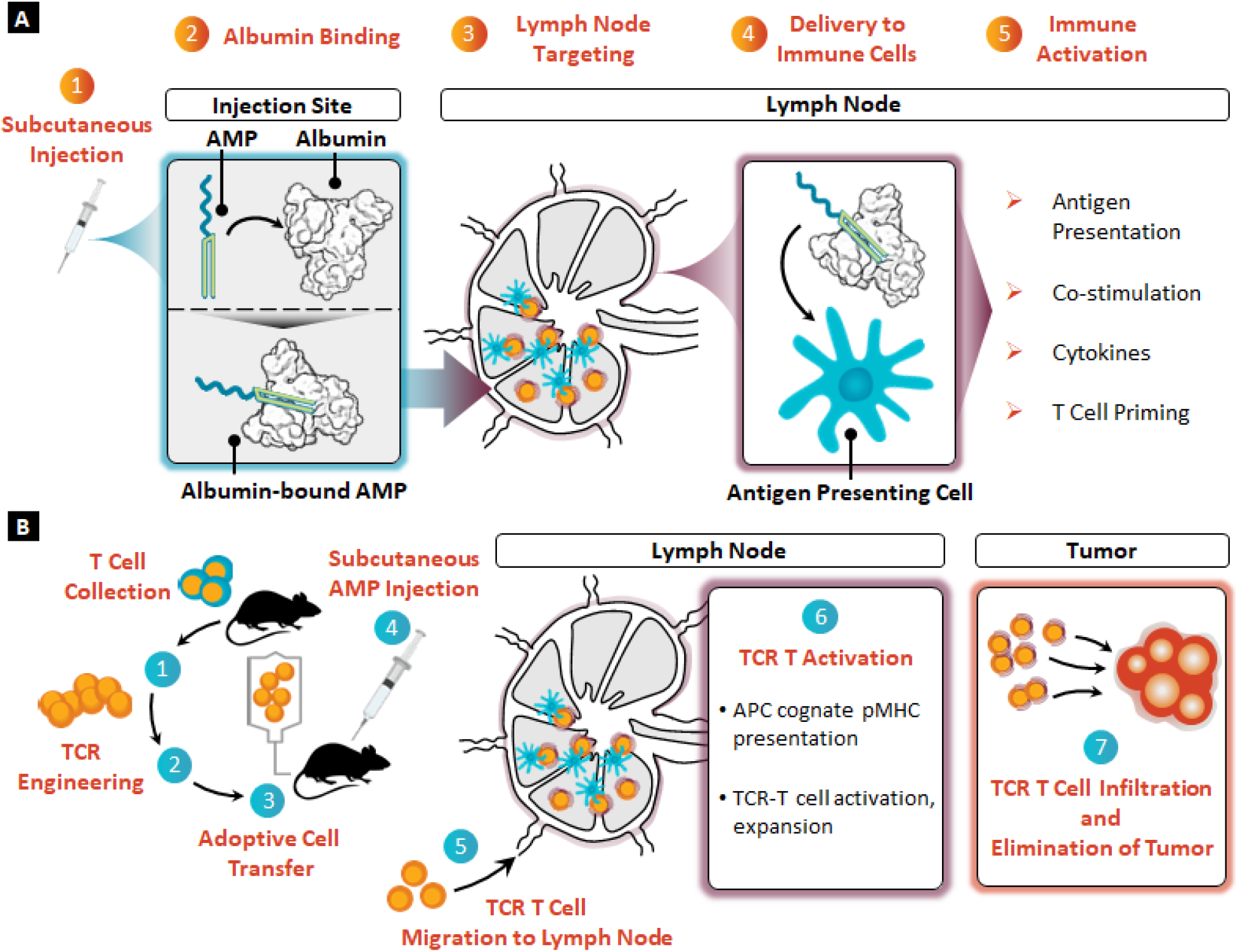
Design of a lymph node-targeted TCR-T cell therapy vaccine combination. **(A)** Schematic depicting the lymph node trafficking and immunostimulatory effects of AMP-conjugated vaccine components following administration. AMP-conjugated vaccines are (1) administered subcutaneously before (2) binding to tissue-resident albumin at the injection site. Albumin-bound AMP-conjugates then (3) traffic to the lymph node where (4) they are delivered to immune cells including APCs to (5) stimulate immune mechanisms supporting T cell activation. **(B)** Schematic illustrating the proposed combination of AMP-vaccination with TCR-T cell therapy in lymph nodes, leading to enhanced T cell tumor infiltration and tumor regression. Autologous T cells can be (1) collected, (2) genetically engineered to express TCR constructs of interest, and (3) adoptively transferred intravenously into tumor-bearing mice alongside (4) subcutaneous AMP-vaccination. TCR-T cells then (5) migrate to lymph nodes, where they encounter primed, activated APCs leading to (6) TCR-T cell activation and expansion. Activated TCR-T cells can then traffic to the tumor site where they exhibit (7) enhanced infiltration and tumor elimination.

Given the efficacy of AMP-vaccines to potently expand endogenous effector T cell responses, we theorized that a similar approach could be designed to enhance TCR-T cell therapy against established solid tumors in vivo. Using this approach, conventional collection, activation, genetic programming, and re-infusion of tumor-specific TCR-T cells can be combined with AMP-vaccination including matched cognate-peptide and adjuvant (Fig. 1B). Following administration, TCR-T cells migrating through the lymphatics can colocalize in the lymph node with a population of highly costimulatory, cognate-peptide-presenting APCs in the context of a supportive immune microenvironment unique to these anatomical sites. Through AMP-vaccination, concerted delivery of TCR stimulus with endogenous co-stimulation and cytokine support can effectively promote TCR-T cell activation, expansion, and on-demand reinforcement of anti-tumor effector function in vivo. This also allows establishment of a supportive lymphatic niche for induction of antigen-spreading to diversify immune recognition of tumor-specific targets to promote durable complete remission and long-term protection. Herein, we describe evidence utilizing multiple TCR-T cell models to demonstrate the unique enhancement provided by effective lymph-node targeted AMP-vaccination, including effects on APCs to support TCR-T cell activation and expansion in vitro, enhanced tumor infiltration, and eradication of established solid tumors in a syngeneic murine model. Subsequently, we demonstrate the potential for AMP-peptide-driven enhancement of human TCR-T cells specific for clinically relevant tumor targets indicating the potential for this strategy to improve TCR-T therapeutic activity upon clinical translation.

## Results

### AMP-vaccination enhances TCR-T cell therapy and eradicates established solid tumors

AMP-modification of peptide antigens and molecular adjuvants is known to promote enhanced lymphatic delivery and retention through albumin “hitchhiking” leading to potent expansion of cognate endogenous T cell responses (*23, 28, 29*). Motivated by these findings, we sought to design a lymph-node-targeted vaccination regimen for adoptively transferred cognate TCR-T cell therapy to enhance therapeutic efficacy against established solid tumors. Wildtype C57BL6 (WT) mice were implanted subcutaneously with 5×10^5^ B16F10 melanoma tumor cells for 10 days, resulting in tumor sizes of 40-70 mm^3^. Activated, mCherry-transduced, tumor-specific, Pmel-1 TCR-T cells specific for gp100_25-33_ were then administered on day 10 without any prior conditioning or exogenous cytokine supplementation (irradiation pre-conditioning had previously failed to decrease tumor growth or promote overall survival in this tumor model, fig. S1A-B). TCR-T cell therapy was administered alone or in combination with a multi-dose vaccine boosting regimen consisting of cognate gp100 long peptide and CpG adjuvant (Fig. 2A). To determine the importance of lymphatic targeting for effective vaccination of TCR-T cells, unmodified soluble (SOL) peptide and adjuvant were compared to AMP-modified vaccine components. Tumor growth in mice treated with AMP-vaccination in concert with TCR-T cell infusion was significantly delayed or completely eradicated when compared to mice given TCR-T cells alone or in combination with SOL vaccination (Fig. 2B); consequently, AMP-vaccinated mice had significantly enhanced long-term survival without signs of distress, such as visible weight loss (Fig. 2C, fig. S1C). Enhanced anti-tumor efficacy in AMP-combination treated animals was further associated with increased number and persistence of TCR-T cells in the peripheral blood of treated tumor-bearing animals for up to 75 days following adoptive transfer (Fig. 2D, fig. S1D). Importantly, despite the expansion of TCR-T cell numbers, AMP-vaccine combination treatment did not result in significant systemic levels of inflammatory cytokines indicative of potential for cytokine release syndrome (fig. S1E). In another study, earlier TCR-T-AMP-vaccine combination treatment of B16F10 tumors established for only 7 days resulted in improved anti-tumor efficacy while TCR-T cell therapy alone or in combination with SOL comparators were similarly ineffective (fig. S1F-G).

**Fig. 2:**
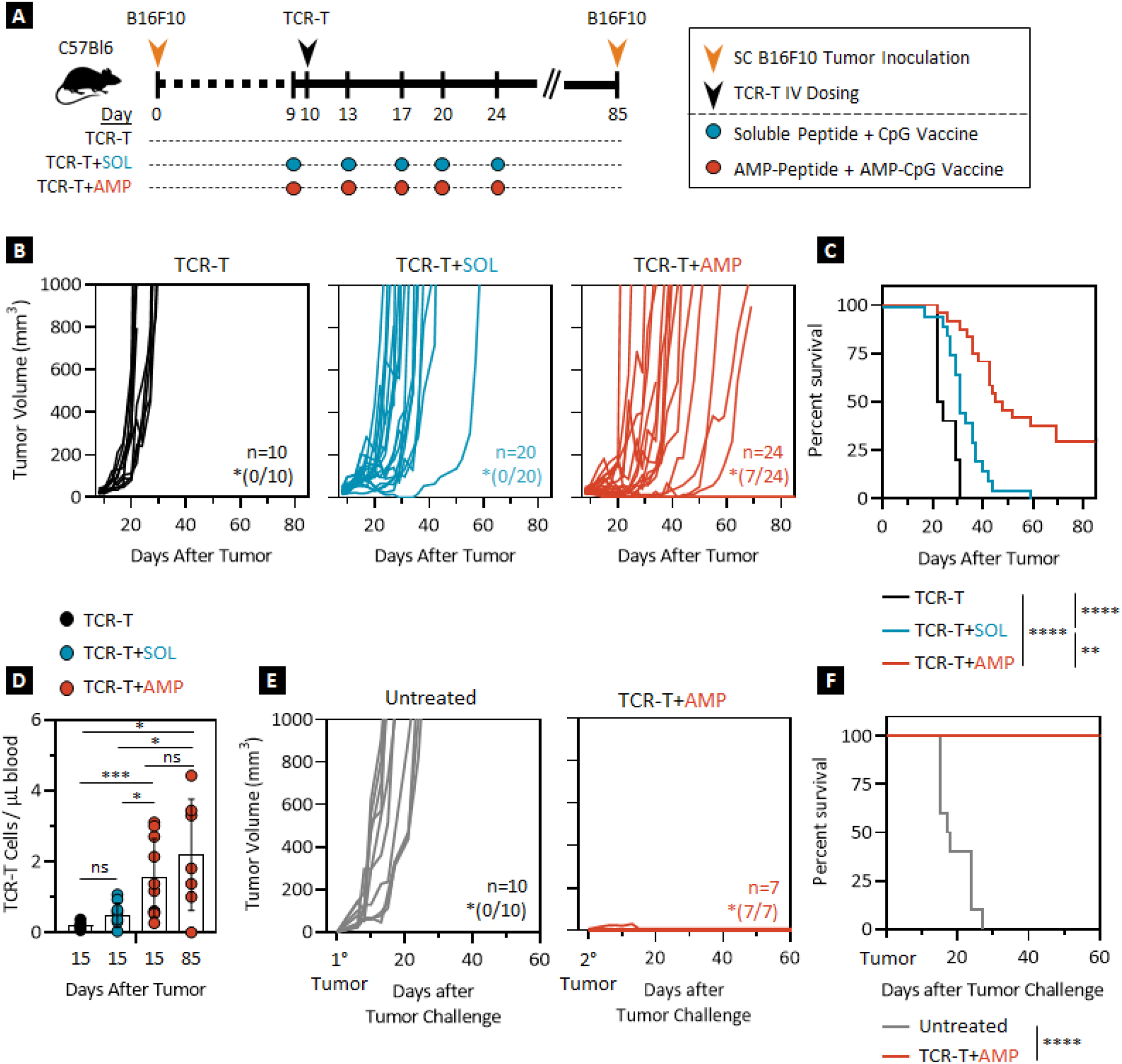
AMP-vaccination enhances TCR-T cell therapy and eradicates established solid tumors. **(A)** Schematic of experimental protocol. **(B)** Individual tumor volume; *indicates number of mice tumor-free at day 85 post tumor. **(C)** Overall survival over time for B16F10 tumor-bearing mice treated with transduced, activated, gp100-specific TCR-T cells alone or in combination with 10 µg peptide and 1 nmol adjuvant in either SOL- or AMP-gp100 vaccine regimens. Data shown are from 3 independent experiments. **(D)** TCR-T cell number per µL of peripheral blood in treated mice at day 15 and 85 after tumor inoculation. Data shown are means ± SDs from 2 independent experiments; p-values were determined by the Mann-Whitney Test. **(E)** Individual tumor volume; *indicates number of mice tumor-free at day 60 post tumor. **(F)** Overall survival for naïve control and surviving TCR-T cell + AMP-vaccine combination treated mice following secondary tumor challenge on day 85. Secondary tumor challenge data shown are from 2 independent experiments. P-values for survival were determined by the log-rank Mantel-Cox Test, with 95% confidence interval. ****, *p* < 0.0001; ***, *p* < 0.001; **, *p* < 0.01; and *, *p* < 0.05.

To evaluate the potential for further protection against relapse, mice previously treated with TCR-T cells and AMP-vaccine that achieved long-term survival were implanted with a secondary B16F10 tumor challenge 75 days after T cell treatment alongside a naïve control group. All long-term surviving mice exhibited continued TCR-T cell persistence (fig. S1H) and rejected growth of the secondary tumor in comparison to the naïve control group in which tumor growth quickly required euthanasia (Fig. 2E-F, fig. S1I-J), confirming that TCR-T cell combination with AMP-vaccination induces durable protection against solid tumor rechallenge in a syngeneic murine model.

### AMP-vaccination enhances TCR-T cell activity *in vitro* and *ex vivo*

Given the enhanced in vivo efficacy of AMP-vaccine combination with TCR-T cell therapy to eradicate advanced solid tumors, we used in vitro and ex vivo analyses to mechanistically understand this combination effect. While in vitro assessment is not expected to fully capture the mechanistic processes critical for activity in vivo, ex vivo analysis provides a useful opportunity to evaluate AMP-mediated contributions to lymph node accumulation and retention, as well as effective uptake, processing, and presentation of cognate peptide by APCs that are in regional proximity to TCR-T cells as they circulate in vivo. In vitro, murine dendritic cells (DCs) were incubated with SOL- or AMP-gp100 peptide prior to culture with previously activated, transduced, and rested cognate TCR-T cells to determine the effect on enhancing TCR-T cell effector function (Fig. 3A). Significantly higher levels of pro-inflammatory interferon gamma (IFNγ) were produced by cultures of TCR-T cells with SOL- or AMP-peptide-pulsed DCs relative to T cells cultured alone or with mock-pulsed DCs (Fig. 3B). SOL- and AMP-peptide pulsed DCs induced increased TCR-T cell expression and co-expression of CD25 and CD69 activation markers compared to mock-pulsed DCs or untreated control T cells (Fig. 3C). Similar enhancements were observed in TCR-T cell cytotoxic and effector function against B16F10 target tumor cells. Here, TCR-T cells stimulated with AMP-pulsed and, to a lesser extent, SOL-pulsed DCs exhibited enhanced specific lysis of B16F10 tumor targets (Fig. 3D) and associated secretion of pro-inflammatory IFNγ (Fig. 3E) in comparison to TCR-T cells left untreated or exposed to mock-pulsed DCs. These data illustrate the ability of both SOL- and AMP-peptides to promote DC-mediated enhancement of TCR-T cell function in vitro. As expected, given the limitation of in vitro assessments, these trends were not fully consistent with the relative enhancement in anti-tumor activity we observed previously for SOL- and AMP-vaccination in vivo.

**Fig. 3:**
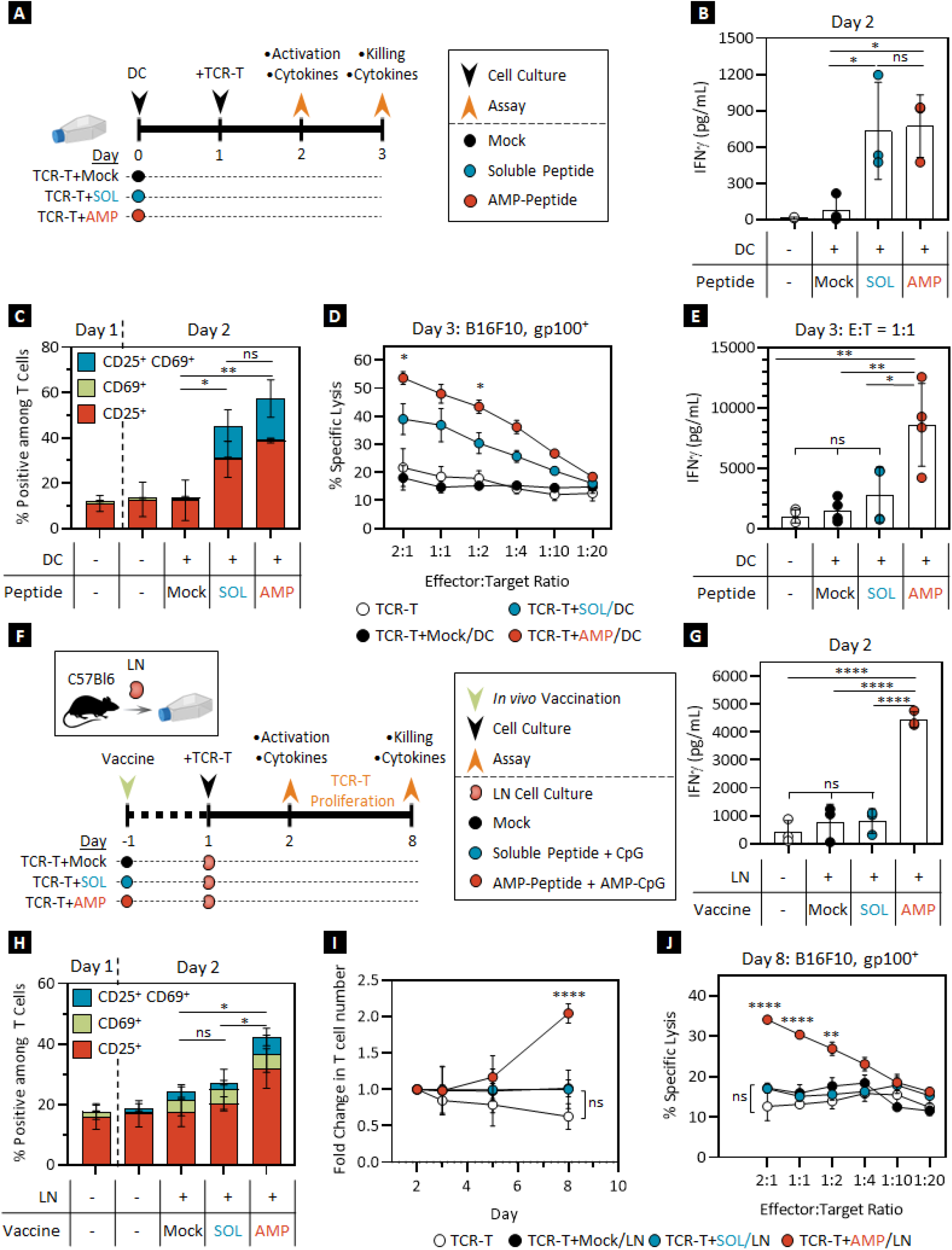
AMP-vaccines differentially enhance TCR-T cell activation and function in vitro and ex vivo. **(A)** Schematic of in vitro experimental protocol. **(B**) IFNγ production determined by Luminex, **(C)** CD25/CD69 expression assessed by flow cytometry, and **(D)** specific tumor lysis in a 24-hour luciferase assay of gp100-specific TCR-T cells after 24-hour co-culture with SOL- or AMP-gp100 peptide-pulsed DC2.4s. **(E)** IFNγ production determined by Luminex of TCR-T cells from a 1:1 effector-to-target ratio in a 24-hour cytotoxicity assay. **(F)** Schematic of ex vivo experimental protocol. **(G**) IFNγ production assayed by Luminex, **(H)** CD25/CD69 expression by flow cytometry, **(I)** longitudinal fold-change in number of gp100-specific TCR-T cells after 24-hour co-culture with lymph node single cell suspension from mock, SOL-, or AMP-gp100 vaccinated mice. **(J)** Specific tumor lysis in a 24-hour cytotoxicity assay of TCR-T cells co-cultured with lymph node single cell suspensions. LN: lymph node. In vitro and ex vivo data shown are means ± SDs, each from 3 independent experiments; p-values were determined by one- or two-way ANOVA. ****, *p* < 0.0001; **, *p* < 0.01; and *, *p* < 0.05.

Therefore, to better understand the role of AMP-driven lymphatic targeting on TCR-T cell vaccination in vivo, we devised an ex vivo approach in which inguinal lymph nodes were harvested from tumor-naive mice 24 hours after dosing with either mock-, SOL-, or AMP-vaccines including gp100 peptide. Single cell suspensions from harvested lymph nodes were subsequently co-cultured at a 1:1 ratio with previously activated, transduced, and rested TCR-T cells (Fig. 3F) before assessment of TCR-T cell activation, cytotoxicity, and effector function. As observed in prior in vitro assessments, pro-inflammatory cytokine levels including IFNγ and tumor necrosis factor alpha (TNFα) were significantly higher in supernatant of TCR-T cells cultured with lymph node single-cell suspension from AMP-vaccinated mice relative to comparator conditions on both days 2 and 8 (Fig. 3G, fig. S2A-C). However, lymph nodes isolated from SOL-vaccinated mice were inactive, consistent with prior studies demonstrating the poor lymphatic accumulation of soluble peptide immunogens and molecular adjuvants (*23, 28*). T cell activation status and proliferation were also significantly enhanced for TCR-T cells cultured with AMP-vaccinated lymph node single cell suspensions in comparison to TCR-T cells cultured with either mock- or SOL-vaccinated lymph node cells (Fig. 3H-I). Finally, ex vivo culture of lymph node cell suspensions from AMP-vaccinated mice allowed TCR-T cells to execute potent anti-tumor cytotoxicity a week later, with significantly greater specific tumor lysis at various effector-to-target ratios relative to TCR-T cells exposed to mock- or SOL-vaccinated lymph node cells or left untreated (Fig. 3J). Consistent with prior correlations of lymphatic delivery with effective APC acquisition and presentation of antigen to activate cognate T cells, these ex vivo data support our hypothesis that lymph node trafficking by AMP-vaccines is an essential mechanism for promoting multiple axes of TCR-T cell function in vivo. These results provide further evidence to determine why SOL-peptide vaccination, while effective at enhancing TCR-T cell activity in vitro, fails to induce a similar enhancement in TCR-T cell anti-tumor response in vivo.

### AMP-vaccination induces DC activation and inflammation in lymph nodes

Given the stimulatory effect of AMP-vaccinated lymph node single cell suspensions on TCR-T cells ex vivo, we then investigated the effect of AMP-vaccination upon both TCR-T cells and APCs within lymph nodes in vivo to understand the correlation between lymph node activation and anti-tumor activity. As before, WT mice were implanted subcutaneously with B16F10 tumor cells for 10 days before treatment with activated, transduced tumor-specific TCR-T cells alone or with SOL- or AMP-vaccination followed by lymph node collection 7 days later (Fig. 4A). Flow cytometric analysis showed that while SOL-vaccination had no effect, AMP-vaccination induced significant increases in the number of lymph node resident TCR-T cells compared to mock- or SOL-vaccinated animals (Fig. 4B). In addition, CD8^+^ T cells within AMP-vaccinated lymph nodes were more functional in secreting pro-inflammatory cytokines in response to a tumor-specific antigen after stimulation ex vivo with gp100 peptide (Fig. 4C). These findings were consistent with our prior observations of TCR-T cells stimulated ex vivo with AMP-vaccinated lymph node single cell suspensions and the known relative lymphatic targeting efficiency of SOL-versus AMP-immunogens, demonstrating that AMP-vaccination has similar effects upon adoptively transferred T cells in vivo.

**Fig. 4:**
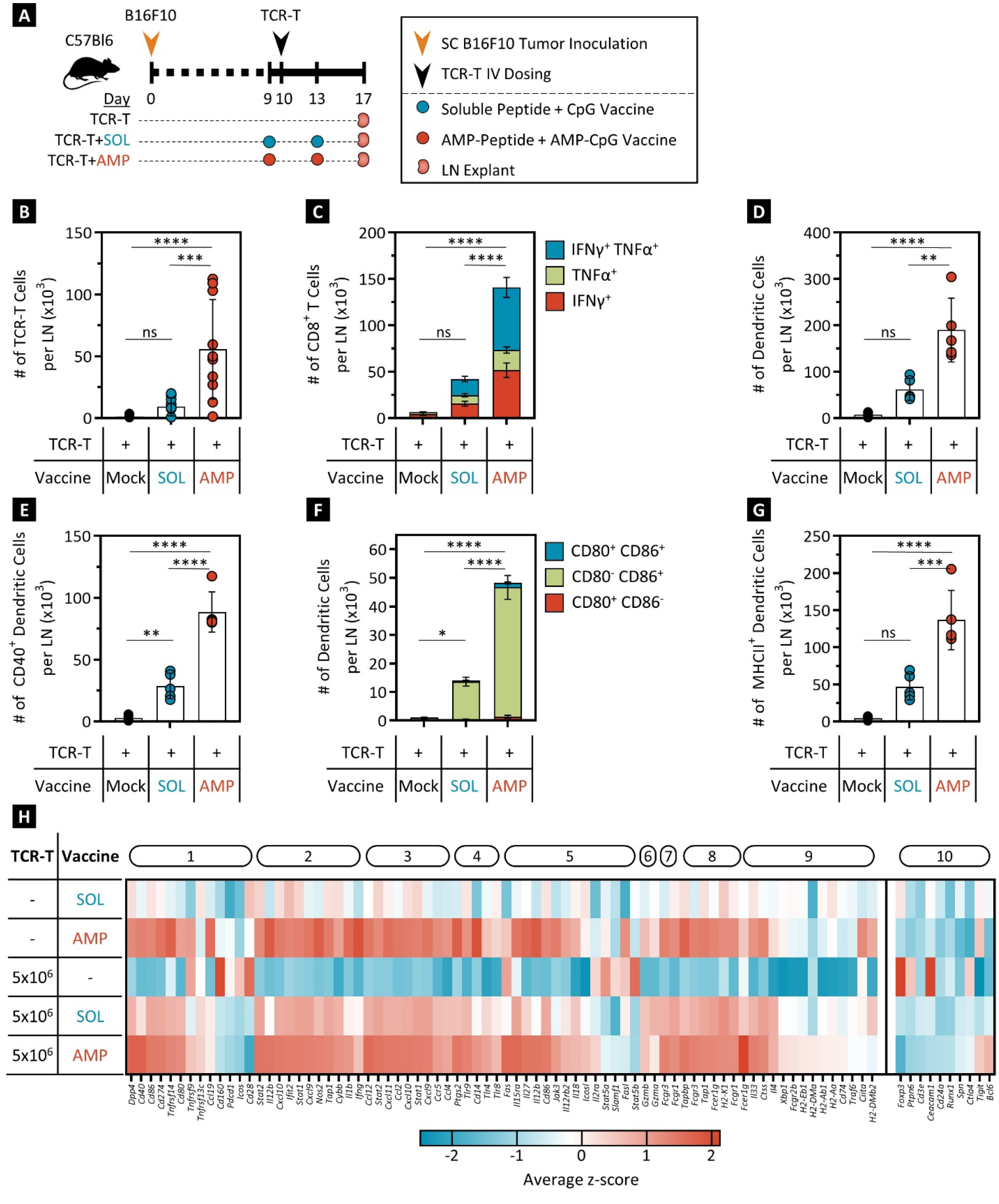
AMP-vaccination induces DC activation and inflammatory transcriptional reprogramming in draining lymph nodes. **(A)** Schematic of experimental protocol. **(B)** Number of gp100-specific TCR-T cells within inguinal lymph nodes. **(C)** Pro-inflammatory cytokine production of CD8^+^ T cells isolated from inguinal lymph nodes when stimulated with gp100 peptides ex vivo as determined by flow cytometry. **(D)** CD11b/c^+^ DC numbers found within inguinal lymph nodes following gp100-specific TCR-T cell therapy combination with mock, SOL-, or AMP-gp100 vaccination assessed by flow cytometry. **(E)** CD40, **(F)** CD80/CD86, and **(G)** MHC Class II expression on DCs in the lymph nodes of treated mice as determined by flow cytometry. **(H)** Heatmap representation of clustered, differentially expressed genes in lymph nodes among the various treatment groups. Annotations describing the biological processes associated with the clustered genes can be found in fig. S3. LN: lymph node. Data shown are means ± SD from 2 independent experiments; p-values were determined by one- or two-way ANOVA. ****, *p* < 0.0001; ***, *p* < 0.001; **, *p* < 0.01; and *,*p* < 0.05.

Previous studies have shown efficient AMP-peptide and adjuvant trafficking to lymph nodes (*23, 30*), and we hypothesized that concerted activation and antigen presentation by APCs in the lymph nodes might be a critical mechanism contributing to the observed improvements in lymphatic induction of T cell expansion and effector function acquisition. Indeed, we observed that AMP-vaccination induced significant increases in the total number of CD11c^+^ DCs, as well as the number of activated DCs expressing costimulatory or antigen-presentation markers CD40^+^, CD80/CD86^+^, or MHC Class II^+^ in lymph nodes 1 week after initiating TCR-T cell therapy (Fig. 4D-G). By comparison, lymph nodes collected from SOL-vaccinated mice exhibited elevated but significantly lower numbers of total DCs and DCs displaying various markers of activation. These findings correlate the potent induction of lymph node DC activation to our observations of elevated TCR-T cell numbers and function demonstrating the critical importance of effective lymphatic adjuvant targeting for enabling the activity of epitope vaccination regimens to enhance adoptive TCR-T cell therapy.

To more broadly assess the differential activation status of the lymph node compartment and the relative contribution of TCR-T cell therapy, SOL- or AMP-vaccination, or their combination, RNA was extracted from lymph nodes collected as before and analyzed by a 561-gene nCounter Mouse Immunology panel to perform an unbiased analysis of the lymph node transcriptome. Z-scores were calculated for the data set and genes were clustered and plotted based upon their gene ontology classification (Fig. 4H, fig. S3). Compared to SOL-vaccination with and without TCR-T cell infusion, AMP-vaccination elevated numerous gene transcripts for pathways associated with inflammation, immune activation, and antigen presentation, while avoiding gene signatures associated with T cell anergy or dysfunction. Upregulated transcripts included co-stimulatory molecules *Cd40* and *Cd86*, inflammatory *Il12b*, *Ifng*, and *Gzmb*, as well as *Tap1* and *Tapbp* which are associated with antigen presentation. Importantly, the elevated relative expression of these pro-inflammatory genes did not correlate with an upregulation of transcripts or pathways commonly implicated in immune regulatory mechanisms, T cell anergy, or exhaustion, including *FoxP3*, *Ctla4*, and *Ceacam1*. Together these data reinforce our observations that AMP-vaccination is effective for inducing numerous and comprehensive immune mechanisms important for supporting and enhancing T cell priming within lymph nodes.

### AMP-vaccination enhances the infiltration and function of TCR-T cells in the tumor microenvironment and leads to epitope spreading against diverse tumor targets

We have shown that AMP-vaccination enhances anti-tumor responses in TCR-T cell-treated tumor-bearing mice, and this effect is correlated with improved in vivo lymph node activation mediated by efficient AMP-peptide and adjuvant accumulation in draining lymph nodes in contrast to soluble comparators (*23, 28*). We further sought to assess the connection between AMP-vaccination induced lymph node responses and anti-tumor immune activity within the tumor itself by observing characteristics of adoptively transferred TCR-T cells and endogenous T cells isolated from the tumor parenchyma. As before, we established B16F10 tumors in mice for 10 days before treatment with activated, transduced, tumor-specific TCR-T cells and either a mock-, SOL-, or AMP-vaccine combination regimen before sacrifice and excision of tumors 7 days later (Fig. 5A). Flow cytometric analysis of single cell suspensions generated from the tumor samples demonstrated that tumors from mice that received TCR-T cell therapy alongside an AMP-vaccination regimen had significantly higher infiltration of adoptively transferred TCR-T cells in comparison to mice given a mock- or SOL-vaccination regimen (Fig. 5B). In addition, total tumor-infiltrating T cells (including endogenous and transferred) from AMP-vaccinated animals were more activated, proliferative, and functional as illustrated by CD25, Ki67, or IFNγ and granzyme B expression (Fig. 5C-E). Despite the prior observation that SOL-vaccination can drive moderate improvements in lymph node DC numbers and activation status, these did not translate into appreciable improvements in tumor-infiltrating T cell responses compared to TCR-T cell therapy alone. Notably, the effective induction of enhanced infiltration, and overall fitness of tumoral T cells by AMP-vaccine combination was not associated with over-activation or exhaustion of T cells. While we observed upregulation of the PD-1 activation marker, this was not accompanied by increases in PD-1/TIM-3/LAG-3 double or triple positive T cells that would indicate T cell dysfunction (fig. S4A).

**Fig. 5:**
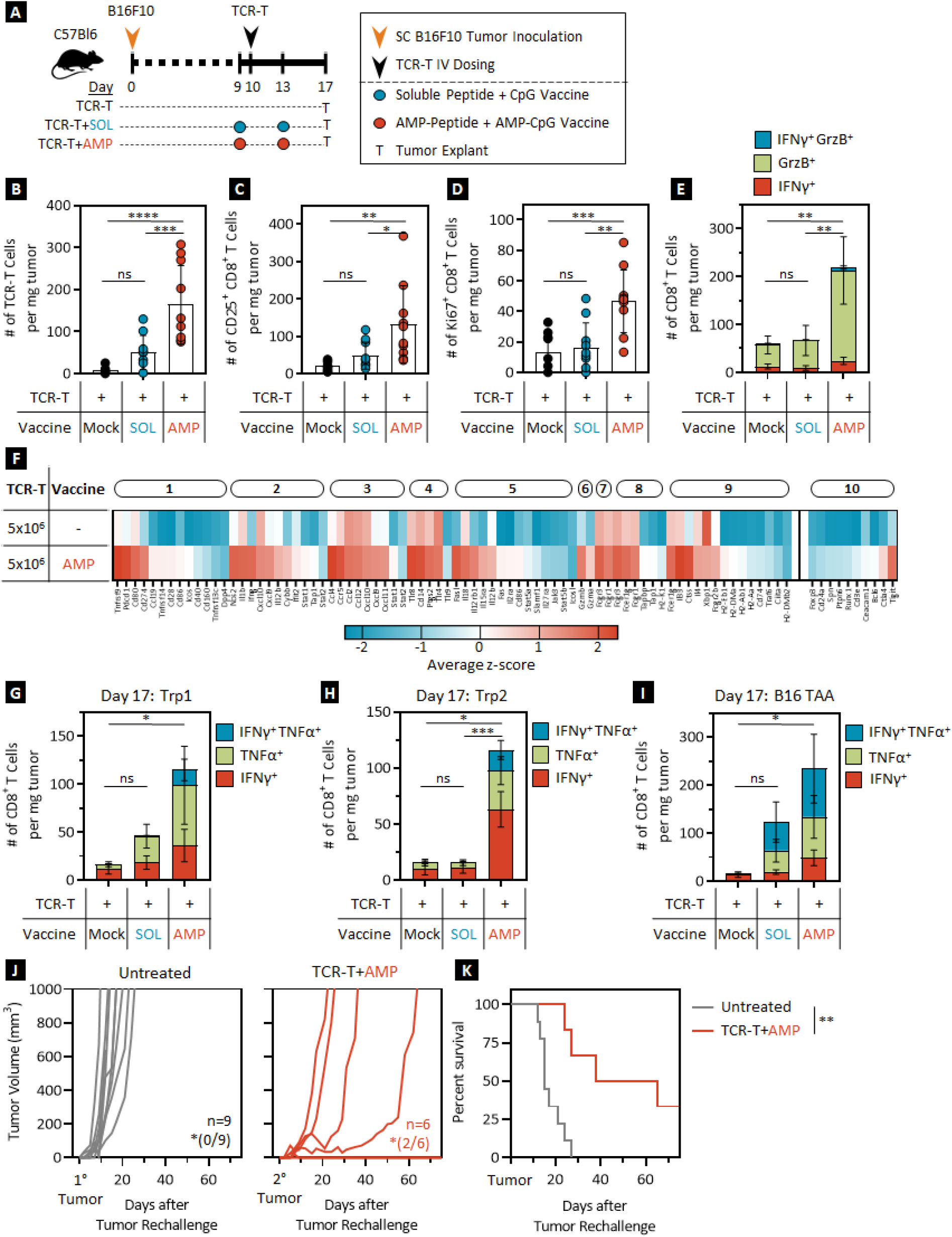
AMP-vaccination enhances the infiltration of activated and proliferating TCR-T cells while inducing inflammatory transcriptional reprogramming, and epitope spreading in the tumor microenvironment. **(A)** Schematic of experimental protocol. **(B)** Accumulation of gp100-specific TCR-T cells within tumor samples from treated mice determined by flow cytometry. Number of **(C)** activated CD25^+^, **(D)** proliferating Ki67^+^, and **(E)** IFNγ- and GrzB-secreting CD8^+^ T cells found within tumors of treated mice assessed by flow cytometry. **(F)** Heatmap representation of clustered differentially expressed genes in tumors from the various treatment groups. **(G-I)** IFNγ and TNFα cytokine secretion detected by flow cytometry from T cells excised from tumor samples of mock, SOL-, or AMP-gp100 vaccinated and gp100-specific TCR-T cell treated tumor-bearing mice following ex vivo stimulation with non-targeted T cell epitopes including **(G)** Trp1, **(H)** Trp2, and **(I)** a panel of B16F10 antigens (M30, Tyrp-1, P15E, M27, M47, M48). **(J)** Individual tumor volume; *indicates number of mice tumor-free at day 75 post tumor. **(K)** Overall survival for naïve control and surviving TCR-T cell + AMP-gp100 vaccine combination treated mice following anti-Thy1.1 depletion and secondary tumor challenge on day 85. Secondary tumor challenge data shown are from 2 independent experiments. Data shown are means ± SDs from 2 independent experiments; p-values were determined by the Mann-Whitney test, one-, or two-way ANOVA. P-values for survival were determined by the log-rank Mantel-Cox Test, with 95% confidence interval. ****, *p* < 0.0001; ***, *p* < 0.001; **, *p* < 0.01; and *, *p* < 0.05.

We then conducted an unbiased transcriptome analysis of the tumor microenvironment to more broadly examine changes in the immune profile associated with TCR-T cell therapy alone versus in combination with AMP-vaccination (Fig. 5F, fig. S3). Relative to tumors from animals treated with TCR-T cell monotherapy, tumor samples isolated from mice treated with the TCR-T cell, AMP-vaccine combination exhibited upregulation in numerous gene transcripts that are instrumental in multiple axes of immune cell activation. Specifically, AMP-vaccination led to upregulation of transcripts associated with T cell activation, interferon signaling, T cell proliferation, and MHC presentation. For example, transcripts related to T cell activation such as *Tnfrsf9* (4-1BB) and *Cd80*, effector T cell homing chemokines such as *Cxcl10* and *Cxcl9*, and T cell proliferation such as *Il18* or *Il15Ra* were notably upregulated in response to AMP-vaccine combination therapy. As in the associated lymph node analyses, we found that upregulation of these various immune activating and inflammatory transcripts, including *Pdcd1* (PD-1), was not accompanied by increases in regulatory transcripts or processes associated with T cell anergy, dysfunction, or tumor promotion such as *FoxP3*, *Ceacam1*, or *Bcl6*. Considered together with the observation of AMP-vaccine-dependent induction of T cell infiltration and improved fitness, these results indicate a more comprehensive shift in the local tumor-immune-microenvironment associated with AMP-vaccine-driven lymphatic activation.

Though in vivo activation of adoptively transferred TCR-T cells by AMP-vaccination is a critical factor in driving anti-tumor activity, we hypothesized that AMP-vaccine-induced activation of the lymph nodes might simultaneously promote endogenous T cell responses specific to other untargeted tumor-associated or neoantigen epitopes. To investigate this, tumor samples from the various treatment groups were stimulated with a panel of peptides previously reported to be present in the B16F10 mutanome (*31*), and the presence of cognate T cells was assessed by flow cytometric detection of effector cytokine production. Tumors from mice treated with TCR-T cells combined with AMP-vaccination exhibited significantly more infiltrating, tumor-antigen-specific T cells relative to comparators from TCR-T cell monotherapy or SOL-vaccination groups (Fig. 5G-I). This increase in AMP-vaccine induced tumor infiltration by endogenous tumor-specific T cells suggests a potential role for this mechanism to promote broader, more potent anti-tumor efficacy and durable protection from tumor progression driven by antigen-loss or heterogeneity. To determine the contribution to anti-tumor effect elicited by the licensing of endogenous T cells against tumor-associated- and neo-antigens, mice that had previously overcome a B16F10 subcutaneous tumor through TCR-T cell and AMP-vaccination therapy were depleted of adoptively transferred Thy1.1 TCR-T cells prior to secondary challenge with B16F10 tumor cells (fig. S4B-D). Consistent with prior observations of complete rejection of secondary challenge by host animals with a replete TCR-T memory cell response, a subset of host mice depleted of adoptively transferred TCR-T cells showed complete tumor rejection, however others displayed delayed, yet progressive tumor growth (Fig. 5J-K). This effect illustrates both the contribution of epitope spread within the endogenous T cell repertoire to promoting overall anti-tumor activity and the potential of AMP-vaccination to support this mechanism in a syngeneic murine tumor model.

### AMP-vaccination induces lymphatic DC activation and enhanced mutant KRAS (mKRAS)-specific TCR-T cell responses *in vivo* in WT and HLA expressing mice

To further assess the potential activity and mechanisms associated with AMP-vaccination to enhance TCR-T cells representative of translational candidates for treating human disease, we conducted studies of lymphatic trafficking, APC activation, and activation of mutant KRAS (mKRAS)-specific TCR-T cells in a transgenic mouse strain conducive to TCR-T cell priming through matched human HLA, TCR engagement. We first explored lymphatic targeting in WT mice treated with vaccines including FITC-labelled, SOL- or AMP-mKRAS-G12D peptide immunogens. Inguinal and axillary lymph nodes were harvested 24 hours after treatment and imaged ex vivo for peptide-associated fluorescent signal (Fig. 6A, fig. S5A). AMP-vaccinated animals exhibited enhanced peptide accumulation in local and distal draining lymph nodes highlighting the relative lack of lymph node exposure achieved with conventional soluble peptides, and the ability of AMP-modification to induce robust lymphatic delivery targeting multiple regionally distinct lymph node groups. Flow cytometric analysis of lymph node isolates confirmed and expanded these findings showing significant increases in the fraction of lymph node-resident DCs and macrophages observed to simultaneously acquire FITC-labelled peptide and exhibit markers of APC-activation, CD80 and CD40. By contrast, SOL-vaccinated comparators demonstrated no increase in lymph node APC populations positive for antigen and costimulatory markers (Fig. 6B-E). These data confirm that clinically relevant AMP-peptides traffic to draining lymph nodes more efficiently than conventional soluble peptides. This finding led us to investigate whether these peptides induce TCR-T cell engraftment, activation phenotype, and effector function in an HLA-expressing, clinically-relevant murine system.

**Fig. 6:**
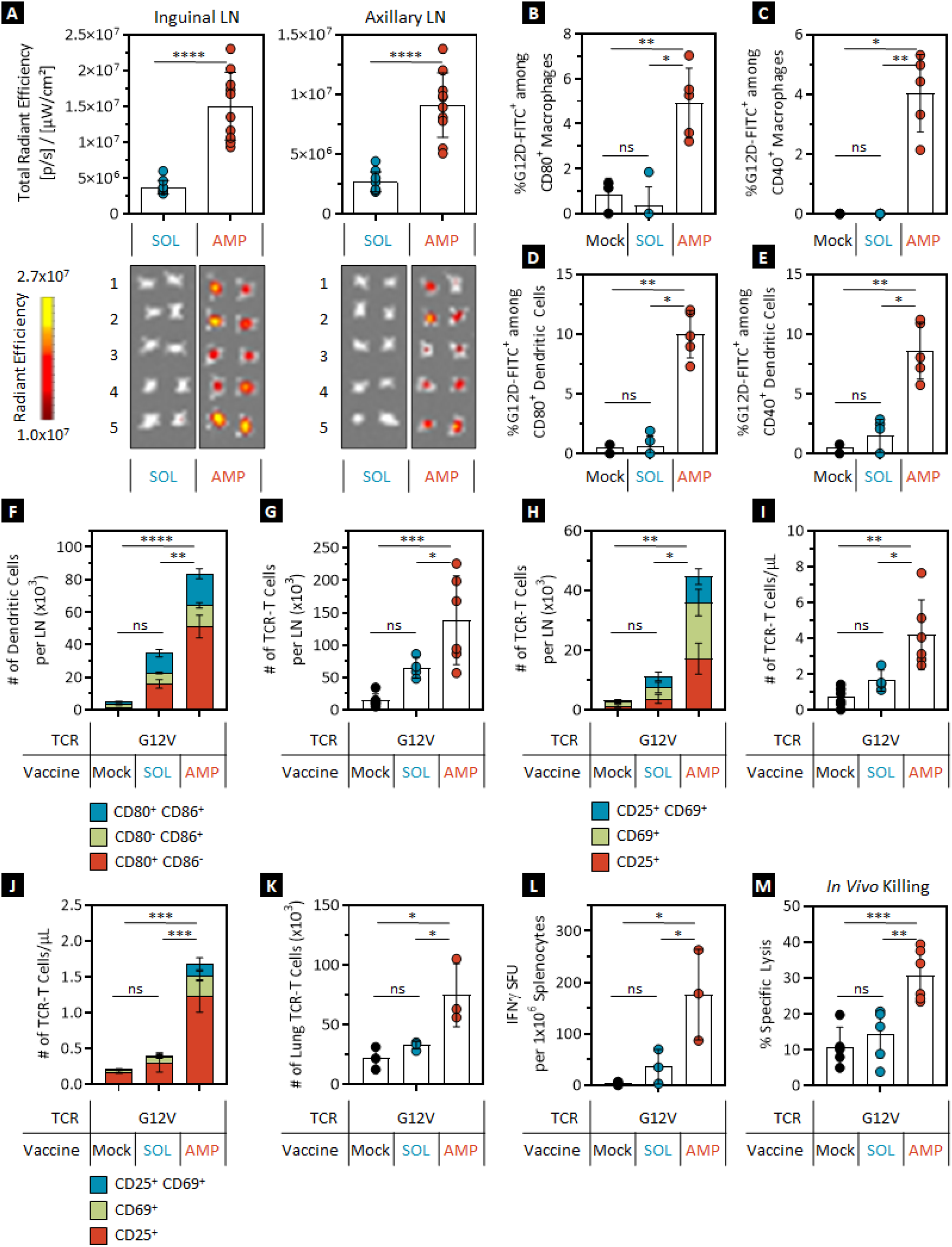
AMP-vaccination enhances the number of activated and functional mKRAS-specific murine TCR-T cells in wild type and A*11:01 transgenic mice in vivo. **(A-E)** Immunization of C57Bl/6J mice with SOL- or AMP-vaccines containing FITC-labeled mKRAS-G12D peptides. **(A)** IVIS imaging and quantification of fluorescent signal in inguinal (left) and axillary (right) lymph nodes 24 hours post injection. Association of FITC-labeled mKRAS-G12D with **(B, C)** CD80^+^ or CD40^+^ macrophages and **(D, E)** CD80^+^ or CD40^+^ DCs 24 hours post injection determined by flow cytometry. **(F-M)** Combination therapy of HLA A*11:01-expressing mice with HLA A*11:01 T cells transduced with an mKRAS-G12D-specific TCR alongside mock-, SOL-, or AMP-mKRAS-G12D peptide vaccination. Enumeration of **(F)** CD11b/c^+^ DCs and **(G, H)** activated mCherry^+^ TCR-T cells within inguinal lymph nodes of treated mice as detected by flow cytometry. Number of CD25/CD69^+^ activated murine mCherry^+^ TCR-T cells **(I, J)** in peripheral blood and the lungs **(K)** of AMP-mKRAS-G12D vaccinated animals in comparison to SOL-vaccinated or mock-treated animals as determined by flow cytometry. **(L)** Splenocyte IFNγ Spot Forming Units (SFUs) following ex vivo stimulation with mKRAS-G12V peptides in an ELISPOT assay and **(M)** TCR-T cell specific lysis of mKRAS-G12V-pulsed splenocytes assessed by flow cytometry in an in vivo killing assay. LN: lymph node. Data shown are means ± SDs from 1 or 2 independent experiments; p-values were determined by the Mann-Whitney test, or one- or two-way ANOVA. ****, *p* < 0.0001; ***, *p* < 0.001; **, *p* < 0.01; and *, *p* < 0.05.

HLA A*11:01^+^ donor murine T cells were transduced with bi-cistronic TCR-mCherry constructs specific for clinically-relevant mKRAS-G12D or -G12V peptide epitopes and infused into naïve HLA A*11:01^+^ host mice in combination with either mock-, SOL-, or AMP-vaccination regimens containing cognate peptide immunogens (fig. S5B). As observed previously in WT mice, only administration of AMP-vaccine led to a significant increase in lymph node APC activation with elevated numbers of CD80/CD86^+^ and MHC Class II^+^ activated DCs observed relative to mock- or SOL-vaccination regimens (Fig. 6F, fig. S5C-D). AMP-vaccination also induced a significant increase in the number of total TCR-T cells and activated CD25^+^/CD69^+^ TCR-T cells within the lymph nodes and the peripheral blood (Fig. 6G-J) consistent with AMP-vaccine induced TCR-T cell priming, expansion and activation in the lymph nodes, supporting enhanced circulating peripheral TCR-T cell populations. Combination of TCR-T cells with AMP-vaccination further induced elevated numbers of TCR-T cells within the lungs, a tissue of high potential clinical interest for TCR-T cell therapy intervention in human mKRAS-driven disease progression (Fig. 6K).

In addition to enhanced persistence and activation of TCR-T cells, combination of TCR-T cell therapy with AMP-vaccination augmented various axes of TCR-T cell effector function. Splenocytes isolated from mice treated with TCR-T cells in concert with AMP-vaccination produced significantly higher numbers of IFNγ-secreting cells relative to mock- or SOL-vaccinated comparators (Fig. 6L). Finally, animals treated with TCR-T cells combined with AMP-vaccination exhibited substantially higher levels of in vivo specific lysis against mKRAS-peptide-pulsed targets in comparison to animals that were given mock- or SOL-vaccination regimens, illustrating the potential of AMP-vaccination to enhance the efficacy of clinically relevant TCR-T cells in vivo (Fig. 6M).

### Clinically relevant human TCR-T cells are activated by AMP-peptide pulsed autologous DCs in vitro

To explore the translational relevance of AMP-vaccination to boost TCR-T cell function, we devised an in vitro system in which peptide-pulsed autologous human DCs were used to stimulate activated, transduced, and rested TCR-T cells across multiple clinically relevant tumor targets and HLA haplotypes. Initially, NY-ESO-1-specific TCRs were transduced into human HLA A*02:01^+^ T cells and, following a rest period, were co-cultured with autologous matured HLA A*02:01^+^ DCs previously pulsed with AMP-NY-ESO-1 peptide or vehicle only (fig. S6A). Exposure to AMP-peptide-pulsed DCs resulted in significant activation of human TCR-T cells in comparison to T cells left untreated or exposed to DCs alone as demonstrated by higher levels of CD25 and CD69 co-expression (Fig. 7A). Consistent with activation marker upregulation, NY-ESO-1-specific TCR-T cells co-cultured with AMP-peptide-pulsed DCs, secreted significantly higher levels of pro-inflammatory IFNγ, TNFα, and IL-2 cytokines in comparison to T cells cultured with unlabeled DCs (Fig. 7B, fig. S6B-C). Longitudinal analysis of total TCR-T cell numbers demonstrated significant AMP-peptide dependent expansion of TCR-T cells with increased memory phenotype, consistent with activation-induced proliferation upon cognate antigen recognition (Fig. 7C, fig. S6D-E). Activated TCR-T cells were harvested following DC co-culture and assessed for cytotoxic activity against an HLA-matched, antigen-positive A375 GFP/LUC tumor cell line. NY-ESO-1-specific TCR-T cells exhibited enhanced specific target lysis following activation with AMP-peptide pulsed DCs indicating improved per-cell cytotoxic potential against targets expressing relevant levels of antigen presentation components and NY-ESO-1 protein (Fig. 7D).

**Fig. 7:**
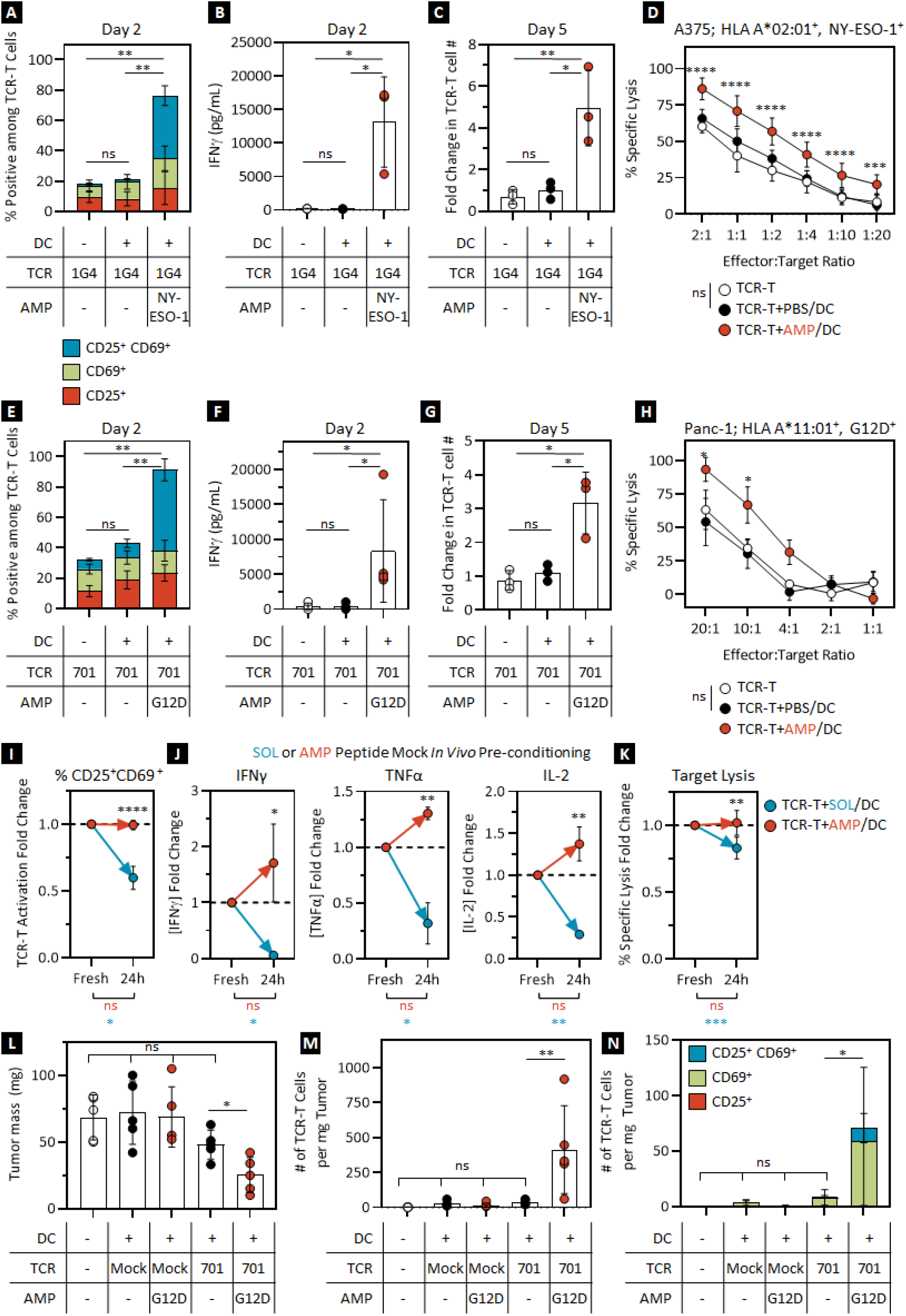
Clinically relevant human TCR-T cells with various specificity are functionally enhanced by AMP-peptide-pulsed autologous DCs in vitro. **(A-D)** NY-ESO-1-specific human TCR-T cells (1G4) were cultured with peptide-pulsed autologous DCs as illustrated in fig. S6A. **(A)** CD25/CD69 expression on TCR-T cells as assessed by flow cytometry, **(B)** production of IFNγ by Luminex, and **(C)** fold-change in TCR-T cell number following co-culture of NY-ESO-1-specific HLA A*02:01 TCR-T cells with mock or AMP-NY-ESO-1 peptide-pulsed autologous human DCs. **(D)** In vitro cytotoxicity of NY-ESO-1-specific TCR-T cells with or without in vitro DC co-culture against HLA-matched, antigen-positive A375 tumor cells in a luciferase killing assay at various effector-to-target ratios. **(E-H)** mKRAS-G12D-specific human TCR-T cells (701) were cultured with peptide-pulsed autologous DCs as illustrated in fig. S6A. TCR-T cell **(E)** CD25/CD69 expression as assessed by flow cytometry, **(F)** production of IFNγ by Luminex, and **(G)** fold-change in TCR-T cell number following co-culture of mKRAS-G12D-specific HLA A*11:01 TCR-T cells with mock or AMP-mKRAS-G12D peptide-pulsed autologous human DCs. **(H)** In vitro cytotoxicity of mKRAS-G12D-specific TCR-T cells with or without in vitro DC co-culture against HLA-matched, antigen-positive Panc-1 tumor cells in a luciferase killing assay at various effector-to-target ratios. **(I-K)** mKRAS-G12D-specific TCR-T cells (701) were cultured with peptide-pulsed autologous HLA A*11:01^+^ DCs previously pulsed with freshly diluted SOL- and AMP-peptides or peptides that were incubated for 24 hours at 37 °C in 10% human serum to mimic in vivo conditions as illustrated in fig. S8A. **(I**) CD25/CD69 expression as assessed by flow cytometry and **(J**) production of IFNγ, TNFα, and IL-2 as determined by Luminex. **(K)** In vitro cytotoxicity in a luciferase killing assay of mKRAS-specific TCR-T cells. **(L-N)** Panc-1 tumor-bearing NSG mice were treated with mKRAS-G12D-specific TCR-T cells (701) after co-culture with mock or AMP-peptide-pulsed autologous HLA A*11:01^+^ DCs as illustrated in fig. S9A. **(L)** Mass of Panc-1 tumors excised from NSG mice on day 35 post treatment. **(M, N)** Number of tumor-infiltrating activated CD25/CD69 expressing TCR-T cells within tumor samples excised from mice treated as determined by flow cytometry. In vitro data were obtained from 2 or 3 independent experiments, and in vivo data were obtained from 1 experiment. Data shown are means ± SD; comparisons between treatment groups shown in black, intra-group comparisons between conditions shown in corresponding colors. P-values were determined by the Mann-Whitney test, one-, or two-way ANOVA. ****, *p* < 0.0001; ***, *p* < 0.001; **, *p* < 0.01; and *, *p* < 0.05.

These results were recapitulated further in in vitro models of HPV16 E7 and mKRAS-G12D in which human HLA A*02:01^+^ T cells or HLA A*11:01^+^ T cells, respectively, were activated and transduced with either HPV16 E7-specific or mKRAS-G12D-specific TCRs and, following a rest period, were co-cultured with autologous HLA A*02:01^+^ or HLA A*11:01^+^ DCs previously pulsed with AMP-HPV16 E7 or AMP-mKRAS-G12D peptides compared to vehicle only (fig. S7A). Once again, we saw that exposure to AMP-peptide-pulsed autologous DCs resulted in upregulation of TCR-T cell activation markers, secretion of inflammatory cytokines, improved TCR-T cell expansion, and enhanced cytotoxic activity (Fig. 7E-H, fig. S7B-G) indicating the versatility of AMP-peptides to enhance TCR-T cell activity independent of antigen-specificity through AMP-modification of TCR-matched minimal or long peptides. This result was also observed for similar models evaluating HLA A*11:01^+^ / mKRAS-G12V-specific and HLA C*08:02^+^ / mKRAS-G12D-specific TCR-T cells with corresponding AMP-mKRAS-peptide-pulsed autologous DCs (fig. S7H-I).

These results clearly demonstrated the stimulatory effect of AMP-peptides on several clinically relevant TCR-T cells in vitro. The in vitro system used, however, does not fully account for the mechanisms known to contribute to robust AMP-vaccine activity in vivo when compared to a SOL-vaccine analog, including lymphatic targeting and retention, and improved resistance to degradation. To investigate the contribution of enhanced in vivo stability to AMP-peptide stimulation of TCR-T cell activation, AMP-peptides and SOL comparators were exposed in vitro to mock in vivo conditions (human serum, 37°C) prior to pulsing autologous DCs for subsequent co-culture with TCR-T cells (fig. S8A) (*30*). Under these conditions, TCR-T cells cultured with AMP-peptide-pulsed DCs demonstrated no detectable decline in ability to induce TCR-T cell activation, cytokine secretion, or specific target lysis after exposure to mock in vivo conditions (Fig. 7I-K). In contrast, the activity of SOL-peptide was significantly reduced or completely ablated by mock in vivo conditions, consistent with the known sensitivity of these agents to passive or enzymatic processes of degradation abundant in tissues. This effect was recapitulated for mKRAS-G12V-peptides and G12V-specific TCR-T cells to show consistent applicability across TCR constructs and peptide epitopes (fig. S8B-C). Together with prior observations, these results highlight another challenge limiting the potential activity of conventional peptide immunogens and demonstrate a mechanism by which AMP-modification may overcome this limitation to enhance TCR-T cell priming and effector function in vivo.

To investigate whether the in vitro enhancement conferred to clinically relevant mKRAS-specific TCR-T cells led to enhanced in vivo anti-tumor effect, we first co-cultured human TCR-T cells with AMP-peptide- or mock-pulsed autologous DCs in vitro prior to treating Panc-1 (HLA A*11:01^+^, mKRAS-G12D^+^) tumor-bearing NSG mice (fig. S9A). As NSG mice lack an intact and competent immune system, in particular functioning lymph nodes essential for AMP-vaccine mechanisms of action, we boosted the TCR-T cells in vitro prior to infusion. Ten-day Panc-1 tumor-bearing NSG mice were infused with either mock-transduced or mKRAS-G12D-specific TCR-T cells isolated from overnight co-cultures with mock or AMP-peptide pulsed DCs and followed for tumor progression. On day 35 post tumor implantation, the tumors were excised and analyzed for tumor mass, TCR-T cell infiltration and activation. Only the combination of mKRAS-G12D-specific TCR-T cells with AMP-peptide-pulsed DCs led to a significant relative reduction in endpoint tumor mass (Fig. 7L). Corresponding analysis of tumor samples showed that the TCR-T cells exposed to AMP-peptide pulsed DCs in vitro also exhibited significantly higher tumor infiltration (Fig. 7M), higher levels of T cell activation (Fig. 7N), and a lack of exhaustion phenotype (fig. S9B). These data illustrate that DC-mediated AMP-peptide exposure promotes activation and solid tumor infiltration by clinically relevant TCR-T cells associated with enhanced anti-tumor responses.

## Discussion

Despite modest overall response rates and limited durability, the significant therapeutic potential of TCR-T cell therapy suggests that optimization of these modalities could have wide-ranging therapeutic benefit in a multitude of previously intractable solid tumors (*1–3, 5–9*). Among other correlates, clinical failures of TCR-T cell therapies have been associated with poor TCR-T cell engraftment and persistence which can potentially be attributed to a lack of in vivo T cell activating signals (*10, 17*). We have shown that a combination of TCR-T cell therapy with cognate peptide AMP-vaccination provides substantial enhancement of anti-tumor effect including long-term non-progression and resistance to secondary tumor challenge in a substantial fraction of animals. This correlated with increased numbers, persistence, tumor infiltration, and function of TCR-T cells, consistent with previous preclinical and clinical reports associating functional T cell dynamics with better overall survival (*32*).

Multiple strategies have been developed to enhance the curative potential of TCR-T cell therapies, including genetic modification of T-cell-intrinsic biology beyond re-engineered TCR-specificity, as well as the development of combination regimens including supportive exogenous cytokines or other factors. Pro-inflammatory cytokines or other co-stimulatory molecules such as IL-12, IL-15, IL-18, or CD40 ligand (CD40L) can be included in TCR expression cassettes, generating “armored” TCR-T cells with enhanced survival and anti-tumor capabilities (*33–37*). Alternatively, TCR-T cell engraftment and persistence can be enhanced by exogenous provision of cytokines such as IL-2 or IL-15 in combination regimens or through direct T cell modification with cytokine “backpacks” (*38–40*). While these therapeutic modalities have great promise, challenges and limitations have also been reported. Exogenous cytokine supplementation is a non-specific therapy with potential for unwanted toxicity (*39, 41*), often further complicated in practice by short serum half-lives and inadequate therapeutic indices. Alternatively, constitutive expression of pro-inflammatory molecules by armored TCR-T cells seeks to restrict potential toxicities associated with systemic exposure; however the potential for adverse side effects remains and often requires additional genetic modifications to ensure safety (*42–46*). By contrast, immunization has the benefit of repeat dosing potential for safe, on-demand augmentation of engineered T cells in vivo, providing an opportunity for support of long-term T cell functional persistence (*27, 29, 47–49*). In mice bearing syngeneic tumors, repeat dosing of AMP-vaccines was well tolerated with no observation of overt toxicity in combination with TCR-T cell therapy. Prior evaluation has shown that AMP-directed delivery to lymph nodes can reduce potentially toxic systemic inflammation otherwise observed for potent adjuvants due to primary clearance from the injection site through the blood (*23*). In the setting of TCR-T cell combination, AMP-vaccination provides the potential for potent TCR-T cell activation and expansion but restriction of this process to lymph nodes likely provides an additional measure of safety, preventing widespread exposure to inflammatory cytokines associated with potential toxicity. Finally, combination of TCR-T cell therapy with AMP-vaccination provided significant improvements in anti-tumor efficacy without the need for traditional preconditioning regimens or exogenous cytokine supplementation, suggesting the potential for AMP-vaccination to supplant these elements of standard practice which nonetheless are associated with potential safety concerns and limit clinical practice to patients willing and able to undergo preconditioning therapy (*50, 51*).

Collectively, next-generation genetic engineering strategies or exogenous factor combination regimens have focused on targeted provision of selected single agents with powerful but potentially narrow biological impact. While these strategies have demonstrated promise, they inherently lack the ability to engage the full breadth of activating mechanisms employed during natural T cell priming by APCs. Further, while these strategies may advance the goals of improved TCR-T cell engraftment and functional persistence, they are by nature or design restricted to enhancement of adoptively transferred cells with single or oligo tumor-specificity and thus limited in their ability to address the likely mechanisms for tumor-immune escape. More comprehensive T cell activation can be achieved through cognate peptide vaccination to widely engage the known mechanisms for expanding and enhancing the function of T cells orchestrated by APCs. In vivo, AMP-vaccination induced significant increases in activated lymph node-resident DCs. Exposure to these cells in lymph node cell suspensions ex vivo resulted in significant enhancement of TCR-T cell biology, including activation, proliferation, effector function, and per-cell tumor killing capacity. Parallel benefits were observed in vivo in TCR-T cells observed in blood, lymph nodes, and tumor parenchyma corresponding to enhanced overall anti-tumor effect. Evaluation of lymph node transcriptomics further revealed the breadth of immune modulating effects driven by AMP-vaccination. Here, pronounced enhancement of numerous genes and pathways associated with T cell activation and co-stimulation, as well as antigen-presentation and cytokine signaling indicate the comprehensive suite of pro-inflammatory signaling induced by AMP-vaccination and accessible to TCR-T cells within the optimal lymph node microenvironment.

While significant anti-tumor responses have been observed in clinical studies of TCR-T cell therapy, acquisition of tumor-intrinsic escape mechanisms, such as antigen-loss or the downregulation of antigen-presenting machinery can lead to tumor progression even in patients achieving successful engraftment of adoptively transferred TCR-T cells (*4, 8, 16*). Therefore, strategies capable of improving TCR-T cell engraftment and functional persistence are attractive, but potentially limited if not adopted in parallel with efforts to overcome mechanisms of tumor immune escape. Tumor antigens shed through the course of tumorigenic progression or released in response to therapy are known to accumulate in draining lymph nodes and represent a potentially powerful source of information for directing mutanome-specific adaptive immunity if captured and presented properly by the immune response (*52*). In this setting, vaccination resulting in adjuvant-mediated activation of the endogenous APC compartment offers the opportunity to prime tumor-specific endogenous memory or naïve T cells alongside adoptively transferred cells thus expanding the repertoire of effector cells available for anti-tumor effect. In addition to the effect on the adoptively transferred TCR-T cell population measured in syngeneic tumor bearing mice, we found that AMP-vaccination supported the expansion of cytokine-secreting endogenous tumor-infiltrating T cells specific for non-targeted, but tumor-associated-antigens. Induction of antigen-spreading in mice treated with TCR-T cells and AMP-vaccination was further observed to confer partial protection from secondary tumor rechallenge in mice that were depleted of adoptively transferred T cells. These observations illustrate the benefit of AMP-vaccination-induced antigen spreading to diversify the anti-tumor immune response providing potentially critical protection against tumor immune-escape through mutational antigen-loss or downregulation.

While vaccine combinations offer the promise to address many of the inherent issues thought to limit TCR-T cell therapeutic efficacy, vaccine components including unmodified molecular adjuvants and conventional peptides fail to successfully traffic to and stimulate immune cells in the lymph nodes where APC:T cell interactions are orchestrated (*23, 30*). Without the ability to efficiently access and prime APCs within the lymph node, these vaccines are not able to effectively engage adoptively transferred or endogenous T cells in this potent and uniquely endowed immunological site. By contrast, effective lymph node targeting enables more effective APC:TCR-T cell engagement and lymph node delivery of AMP-adjuvant following immunization has the potential to reprogram resident innate immunity to promote endogenous T cell responses to tumor antigens passively accumulated in tumor draining lymph nodes. Throughout the reported studies, comparison between unmodified (soluble) and AMP-vaccine components has demonstrated the importance of effective lymph node delivery and the inherent deficiency of conventional molecular adjuvants and immunogens for lymph node targeting. Furthermore, consistent with prior reports, the observed sensitivity of soluble peptides to degradation when exposed to simulated in vivo conditions represents a significant additional challenge for their utility in vivo, while AMP-modification imparted complete resistance to the mechanisms contributing to loss of peptide integrity. The combined evidence demonstrates that achieving potent TCR-T cell activation in the lymph nodes though AMP-directed vaccine delivery was essential to improving TCR- and endogenous T cell activity, dependent on comprehensive immunological engagement, resulting in greatly enhanced therapeutic efficacy.

Following evaluation in syngeneic murine systems we demonstrated that this therapeutic modality could be easily adapted and applied to enhance the activity of clinically relevant TCR-T cell therapies. To that end, we utilized a murine HLA transgenic model as well as in vitro and in vivo human TCR-T cell experimental systems to show direct clinical relevance of the combination of AMP-vaccination and TCR-T cell therapy. HLA transgenic mice treated with TCR-T cells combined with AMP-vaccination maintained substantially higher levels of persistent TCR-T cell populations and exhibited enhanced in vivo specific lysis of peptide-pulsed targets, illustrating the potential of AMP-vaccination to enhance the efficacy of clinically relevant TCR-T cells in vivo. AMP-peptides were also shown to enhance a multitude of critical T cell characteristics such as activation, proliferation, pro-inflammatory cytokine secretion, and per-cell specific tumor lytic function in a series of human HLA-matched systems with NY-ESO-1-, mKRAS-, and HPV16 E7-targeting human TCR-T cells, illustrating the flexibility of this approach to enhance TCR-T cells of various specificity. Further, the ability to rapidly design and simply manufacture matched AMP-peptides informed by knowledge of TCR cognate peptide sequences makes application of this approach feasible for clinical evaluations in combination with existing or progressing TCR-T cell therapies without the need for any modification to existing manufacturing processes.

Prior studies of vaccine combination with adoptive cell therapy have indicated the promise of this approach for improving anti-tumor efficacy against solid tumors (*29, 48, 49, 53, 54*). Transduction of endogenous anti-viral T cells with tumor-specific chimeric antigen receptors (CARs) offers the potential to enhance the activity of these cells in vivo through immunization against the orthogonal viral antigen (*48, 49, 53*). While these approaches strongly confirm the potential for vaccination to potently enhance tumor-redirected T cell therapy, they nonetheless are dependent on complicated and bespoke multi-cistronic genetic reprogramming for each potential new application. This would be compounded further if applied to TCR-T cell therapy where mispairing of tumor- and viral-specific TCR subunits would present a challenge and dual HLA-restrictions may limit accessible patients. Alternatively, direct vaccine-boosting of CAR-T cells through the chimeric receptor reported by multiple groups offers an effective and streamlined approach (*29, 54*). Notably, in these studies, vaccine-induced activity of endogenous APCs in secondary lymphoid tissues was correlated to enhanced CAR-T cell activity, as well as induction of endogenous antigen spreading responses, consistent with our observations demonstrating the importance of lymph node APCs for enhancing TCR- and endogenous T cell activity. Considered together, the prior evidence for the potential of vaccine combinations with T cell therapy, the reported importance of effective lymphoid APC involvement, and the relative practical advantages of vaccination directly targeting the tumor-specific TCR suggest that AMP-vaccination is a promising combination strategy for TCR-T cell therapy.

In summary, combination of TCR-T cell therapy with AMP-vaccination promotes comprehensive lymph node immune activation to enhance TCR-T cell activity and promote endogenous anti-tumor T cell responses resulting in improved solid tumor eradication and durable protection against recurrence. A clinical trial (NCT04853017) evaluating an mKRAS-specific AMP-vaccine is ongoing, setting the stage for future clinical applications of the AMP lymph node targeting technology. Simple AMP-modification of TCR-cognate peptide sequences provides an attractive potential for broad and rapid application to clinical TCR-T cell programs to potentially enhance TCR-T cell function and promote durable and comprehensive anti-tumor immune responses.

## Materials and Methods

### Vaccine Components

Immunogenic peptides containing cognate minimal TCR epitopes were designed with ectopic terminal cysteines where needed to facilitate chemical modification. Peptides were prepared by solid-phase synthesis, cleaved, and purified using reverse-phase chromatography. Purified peptides were characterized by HPLC analysis and mass spectrometry before conjugation as previously reported (*23, 30*) with DSPE-PEG-Maleimide or N-methylmaleimide to generate the desired AMP-peptides and N-methyl-succinimide (NMS) capped soluble analogs, respectively. The resulting AMP-peptides and soluble NMS-capped peptides were analyzed by HPLC and mass spectrometry to confirm purity (>95%) and identity. The amino acid sequences of the peptides were as follows:

AMP-gp100 C(AMP)AVGALEGPRNQDWLGVPRQL;

SOL gp100 C(NMS)AVGALEGPRNQDWLGVPRQL;

AMP-mKRAS-G12D, C(AMP)YKLVVVGADGVGKSALTI;

SOL mKRAS-G12D, C(NMS)YKLVVVGADGVGKSALTI;

AMP-mKRAS-G12V, C(AMP)YKLVVVGAVGVGKSALTI;

SOL mKRAS-G12V, C(NMS)YKLVVVGAVGVGKSALTI;

AMP-NY-ESO-1, SLLM WITQC(AMP);

### AMP-HPV16 E7, C(AMP)TLHEYMLDLQPETTDLY

Soluble CpG 1826 (SOL-CpG) was obtained from InvivoGen (Catalog no. TLRL1826) and AMP-CpG 1826 (AMP-CpG) was synthesized as previously reported (*23, 30*).

gp100-peptide vaccines consisted of 10 μg of gp100 peptide in its non-conjugated, soluble (SOL-gp100) or Amphiphile-conjugated (AMP-gp100) form combined with 1 nmol SOL-CpG or AMP-CpG, respectively. Mock treatment groups were treated with phosphate buffered saline (PBS) alone.

mKRAS-peptide vaccines consisted of 20 μg of mKRAS-G12D or -G12V peptides in soluble (SOL-mKRAS-G12D or SOL-mKRAS-G12V) or Amphiphile-conjugated (AMP-mKRAS-G12D or AMP-mKRAS-G12V) form combined with 5 nmol SOL-CpG or AMP-CpG, respectively. Mock treatment groups were treated with PBS alone.

#### Animals

Female, 6- to 8-week-old C57BL/6J, B6.Cg-Thy1^a^/CyTg(TcraTcrb)8Rest/J (Pmel-1), and NOD.Cg-*Prkdc^scid^ Il2rg^tm1Wjl^*/SzJ (NSG) mice were purchased from the Jackson Laboratory (Bar Harbor, ME). Female, 6- to 8-week-old CB6F1-Tg (HLA-A*1101/H2-Kb) A11.01 (HLA A*11:01) were purchased from Taconic (Germantown, NY). All animal studies were carried out under an institute-approved Institutional Animal Care and Use Committee (IACUC) protocol following federal, state, and local guidelines for the care and use of animals.

Mice challenged with tumor were measured with calipers to confirm equal tumor load and randomized to different treatment groups 1 day before treatment. Mice were euthanized when tumor growth led to a volume greater than 1,000 mm^3^ calculated by 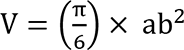 where a is the longest length and b is the shortest length. The investigator was blinded when assessing the outcome. Mice receiving vaccinations were injected with the indicated concentrations of SOL-gp100, AMP-gp100, SOL-mKRAS-G12V, AMP-mKRAS-G12V, SOL-mKRAS-G12D, or AMP-mKRAS-G12D in combination with SOL-CpG or AMP-CpG admixed with PBS. Injections (100 μl) were administered subcutaneously at the base of the tail (50 μl bilaterally) on indicated days. Whole blood samples were obtained utilizing retro-orbital collection method on indicated days. Inguinal lymph nodes, spleens, and tumors were collected at indicated endpoints for downstream analysis.

### Cell Lines

Phoenix-Ecotropic (ECO) packaging cells (ATCC, catalog no. CRL-3214) and Phoenix-Amphotropic (AMPHO) (ATCC, catalog no. CRL-3213) were maintained in DMEM supplemented with 10% heat-inactivated fetal bovine serum (FBS), nonessential amino acids, 2 mmol/L L-alanine-L-glutamine, 4.5 g/L D-glucose, 110 mg/L sodium pyruvate, and 1% penicillin/streptomycin. Murine B16F10 (gp100^+^) cells expressing GFP-firefly luciferase (GFP/LUC) were purchased from Imanis Life Sciences (catalog no. CL068). Cos-7 cell lines expressing HLA A*11:01 and either mKRAS-G12V or mKRAS-G12D, kindly provided by Dr. James Yang from the National Cancer Institute, were modified to express GFP/LUC for killing assays and cultured in DMEM as described above. DC2.4 mouse dendritic cells purchased from Millipore Sigma (Catalog no. SCC142) were maintained in RPMI supplemented with 10% heat-inactivated FBS, nonessential amino acids, 25 mmol/L HEPES, 2 mmol/L L-alanine-L-glutamine, and 1% penicillin/streptomycin. Human Panc-1 (HLA A*11:01^+^, mKRAS-G12D^+^) GFP/LUC cells (GenTarget Inc., catalog no. SC068-LG) were maintained in DMEM as described above. Human A375 (HLA A*02:01^+^, NY-ESO-1^+^) GFP/LUC cells (Imanis Life Sciences, catalog no. CL085) were maintained in DMEM as described above. Human Ca Ski (HLA A*02:01^+^, HPV16 E7^+^) tumor cells (ATCC, Catalog no. CRM-CRL-1550) were modified to express GFP/LUC and maintained in RPMI as described above. Specified cell lines were modified to express GFP/LUC by retroviral transduction.

### Retroviral Constructs, Cell Isolation, and Retroviral Transduction

Murine gp100-specific TCR-T cells expressing mCherry were generated by transducing pmel-1 T cells with plasmids encoding the mCherry construct in the MSGV-retroviral vector. Retroviral vectors were ordered from GenScript and used to transfect Phoenix Eco cells using the Lipofectamine™ 3000 Transfection Reagent and protocol (ThermoFisher, catalog no. L3000075). Pmel-1 or HLA A*11:01 mice were euthanized, and their spleens were harvested. Following mechanical tissue dissociation and red blood cell lysis with (Quality Biological Catalog number 118-156-101CS), T cells were isolated using EasySep protocol (STEMCELL, Catalog no. 19851) and activated with Dynabeads (Gibco, catalog no. 11453D). Cells were expanded in vitro by culturing in T Cell media (RPMI1640 supplemented with 10% heat-inactivated FBS, nonessential amino acids, 1 mmol/L sodium pyruvate, 10 mmol/L HEPES, 2 mmol/L L-glutamine, 1% penicillin/streptomycin, 11 mmol/L glucose, 2 μmol/L 2-mercaptoethanol, and 50 IU/mL of recombinant human IL-2 for human T cell applications and 100 IU/mL for murine T cell applications; Peprotech, catalog no. 200-02). 24 and 48 hours after initial expansion, mouse T cells were spinoculated on RetroNectin-coated plates with viral supernatant collected from Phoenix-ECO (*33*).

To generate the HPV16 E7, NY-ESO-1, and MART-1 specific human TCR-T cells, plasmids including mCherry and/or the specified TCR alpha and beta chains as described in Jin, et al (HPV16 E7) and Drakes et al (NY-ESO-1, 1G4 and MART-1, DMF5) were ordered from GenScript in the MSGV-retroviral vector (*33, 55*). Plasmids encoding the mKRAS-G12D (701, 4095) and mKRAS-G12V (700) targeted TCRs were kindly provided by Dr. James Yang at the National Cancer Institute, Surgery Branch. Phoenix AMPHO cells were transfected with MSGV retroviral vectors using the Lipofectamine™ 3000 Transfection Reagent and protocol (ThermoFisher, catalog no. L3000075). Peripheral blood mononuclear cells (PBMCs) were isolated from leukopaks (STEMEXPRESS) from either HLA A*11:01, HLA A*02:01, or HLA C*08:02 donors. Human T cells were isolated using EasySep Human Kit (STEMCELL, Catalog no. 17951), and cultured in RPMI1640 supplemented with 10% heat-inactivated FBS, 2 mmol/L L-glutamine, and 1% penicillin/streptomycin for 1 day prior to transduction and supplemented with 50 IU/mL of IL-2. Human T cells were activated with human CD3/CD28 Dynabeads (Invitrogen), at a bead:cell ratio of 2:1 and transduced using supernatant collected from Phoenix-AMPHO cells. 24 and 48 hours after isolation, murine and human T cells were transduced by centrifugation on RetroNectin-coated plates. Autologous human DCs were isolated and matured in parallel of human TCR-T cell transduction using a Monocyte EasySep kit and then matured with differentiation and maturation kits (STEMCELL, Catalog nos. 19359, 10988, 10989) prior to antigen presentation to human TCR-T cells.

### In vitro AMP-activation of TCR-T Cells

For murine studies, DC2.4 cells were pulsed in vitro with PBS control or 100 nmol/L of SOL-gp100 or AMP-gp100 for 18 hours. Pulsed DC2.4 cells were then washed 3 times and cultured at a 1:1 ratio with mCherry transduced gp100-specific TCR-T cells that had been previously de-beaded and rested for 3 days to return to a baseline activation status. On days 1 and 7 post culture, supernatant fluid was collected from the cultures for Luminex cytokine analysis and cells were utilized for killing assays against target cells. Additionally, cells were analyzed for activation and phenotype by flow cytometry on days 1, 4, and 7 post co-culture.

For human TCR-T cell AMP-vaccination experiments, mature autologous human DCs were pulsed with PBS or 10 µmol/L of SOL or AMP peptides (mKRAS, HPV16 E7, or NY-ESO-1) for 24 hours. For peptide stability and mock in vivo studies, SOL- or AMP-peptides were incubated at 37°C for 24 hours in 10% human serum containing RPMI prior to DC pulsing. DCs were washed 3 times and cultured at a 1:2 DC:TCR-T cell ratio with mKRAS-, HPV16 E7- or NY-ESO-1-specific human TCR T Cells that had been previously de-beaded and rested for 5 days to return to a baseline activation status. On days 1 and 7 post culture, supernatant fluid was collected from the cultures for Luminex cytokine analysis and cells were utilized for killing assays against target cells. Additionally, cells were analyzed for activation and phenotype by flow cytometry on days 1, 4, and 7 post co-culture.

### In vivo AMP-vaccination of TCR-T Cells

C57/BL6 mice were implanted subcutaneously with 5×10^5^ B16F10 GFP/LUC 10 days prior to adoptive transfer of 5×10^6^ mCherry transduced gp100-specific TCR-T cells. Mice were then vaccinated subcutaneously at the tail base with either PBS (n=10), SOL-gp100/SOL-CpG (n=20) or AMP-gp100/AMP-CpG (n=24) on days −1, 3, 7, 10, 14. Mice were followed for survival, and tumors were measured 3 times per week. To track and assess functionality of circulating TCR-T cells, blood was collected on days 5 and 19 for flow cytometry and ICS analysis. For rechallenge experiments, mice were rechallenged with 5×10^5^ B16F10 GFP-LUC cells on day 75 post T cell transfer with or without intraperitoneal injection of 250 μg anti-Thy1.1 antibody (Days −7, 0, 7, 14, 21, 28, 35) (BioXCell) to deplete adoptively transferred pmel-1 T cells that are Thy1.1 expressing. Depletion of TCR-T cells was confirmed by flow cytometry analysis of processed PBMCs to ensure absence of Thy1.1^+^ cells. For murine mKRAS AMP-vaccination experiments, mKRAS-G12D and -G12V-specific murine TCR-T cells were adoptively transferred to HLA A*11:01 mice. Mice were vaccinated with either PBS, SOL-mKRAS-G12D/SOL-CpG, AMP-mKRAS-G12D/AMP-CpG, SOL-mKRAS-G12V/SOL-CpG, or AMP-mKRAS-G12V/AMP-CpG and circulating TCR-T cells were tracked weekly. End-point harvest analysis was performed by collection and processing of blood, spleen, inguinal lymph nodes, lungs, and tumors (for B16F10-bearing mice) on days 7 or 14 for B16F10 experiments and day 15 for murine HLA A*11:01 mKRAS-G12V and -G12D experiments.

For the human TCR-T cell in vivo tumor model, NSG mice were implanted subcutaneously with 1×10^6^ Panc-1 cells 10 days prior to the TCR-T cell transfer. Human HLA A*11:01 T cells transduced with a mKRAS-G12D-specific TCR or mCherry control construct were activated in vitro by co-culturing with previously AMP-peptide pulsed autologous DCs. TCR-T cells were isolated using a negative T cell isolation kit (STEMCELL, Catalog no. 17951) and injected into the tumor-bearing mice. Tumors were collected 35 days post tumor implantation and analyzed for T cell infiltration, activation, and phenotype.

### Cell Isolation for ex vivo analyses

Tissues (spleen, lymph nodes, lungs, tumor) from vaccinated and TCR-T cell treated mice were excised, mechanically dissociated using a gentleMACS Dissociator (Miltenyi Biotec), then passed through a 70 µm cell strainer. Dissociated cells were centrifuged at 1,500 RPM for 5 minutes at 4°C. The pelleted cells were counted, and an aliquot was resuspended either by cell count (spleen, lymph nodes, and lung) or by mass (tumor) and used for downstream applications. Peripheral blood was collected from study mice at indicated days post T cell treatment and used to assess T cell populations and functionality by flow cytometry. 100 µl peripheral blood was processed in ACK lysis buffer, PBMCs were counted and then resuspended in either 2.5% FBS/PBS prior to antibody staining for flow cytometric analysis of T cell populations or in RPMI1640 medium with 10% FBS, indicated cell stimulation peptide, and 1X Golgi plug (Biolegend) for Intracellular Cytokine Staining (ICS) analysis. The following peptides were used for T cell stimulation in ICS assays: Trp1 (NTPQFENL, GenScript), Trp2 (SVYDFFVWL, GenScript), or a cocktail of B16F10 tumor associated antigens (B16TAA) (M30-VDWENVSPELNSTD, Tyrp-1 175-182-NTPQFENL, p15E 604-611-KSPWFTTL, M27-LCPGNKYEM, M47-GRGHLLGRLAAIVG, M48-DLAVIPAGVVHNWD, all synthesized by GenScript). Cells were transferred to a 96-well U-bottom plate and cultured at 37°C for 18 hours. As a positive control, extra PBMCs from mice receiving TCR-T cells were combined and cultured with 1X Golgi plug and PMA/ionomycin for 18 hours. Cells were pelleted, washed once with PBS, and stained with live/dead violet for 20 minutes in the dark at 4°C. Cells were pelleted again and surface stained for CD3, CD4, CD8, Thy1.1 for 20 minutes on ice. Cells were then washed and permeabilized with the BD Fix/Permeablization kit, according to manufacturer instructions (ThermoFisher) and stained with IFNγ and TNFα antibodies. Cells were washed and resuspended in flow cytometry buffer for analysis immediately or kept at 4°C for analysis on a CytoFLEX flow cytometer the next day.

### Stimulation of TCR-T cells ex vivo with AMP-vaccinated lymph nodes

C57/BL6 mice were injected with mock, SOL-gp100/SOL-CpG, or AMP-gp100/AMP-CpG vaccines; 2 days later, inguinal lymph nodes were collected and mechanically dissociated to generate single cell suspensions. mCherry-transduced gp100-specific TCR-T cells were rested for 3 days prior to culture with lymph node single cell suspensions from either mock, SOL-gp100, or AMP-gp100 vaccinated mice at a 1:1 ratio. On days 1 and 7 post culture, supernatant fluid was collected from the cultures for Luminex cytokine analysis and cells were utilized for killing assays against target cells. Additionally, cells were counted and analyzed for activation and phenotype by flow cytometry on days 1, 4, and 7 post co-culture.

### Cytotoxicity Assay

Mouse or human TCR-T effector cells were plated at various effector to target ratios as specified (20:1, 10:1, 4:1, 2:1, 1:1, 1:2, 1:4, 1:10, or 1:20) against 5×10^5^ antigen-positive target cells expressing GFP/LUC (B16F10, Cos7-mKRAS-G12D/V, Panc-1, CaSki, A375) in 200 µL of total volume in a black-welled 96-well plate. Groups were plated in technical replicates, and additional wells were plated with target only as a background control to determine baseline luminescence. 24 hours after incubation, supernatant was removed and collected for future cytokine analysis. Pelleted cells were incubated with ONE-Glo reagent for 30 minutes at room temperature following the ONE-Glo Luciferase Assay System protocol from Promega (Catalog # E6130). Luminescence was quantified by the Synergy H1 Hybrid reader (BioTek) using the Gen5 Microplate Reader and Imager Software. Specific lysis was determined by analyzing the difference in luminescence of each well relative to the target-only well: % Specific Lysis = 100 – 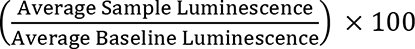

### Luminex cytokine production analysis

Cell culture supernatant or serum from mouse gp100-targeting TCR-T cell experiments were collected and stored at −80°C for future cytokine analysis. The mouse Th17 9-plex Milliplex Map Kit (EMD Millipore, Catalog # MTH17MAG-47k) was used to quantify murine TNFα, IFNγ, GM-CSF, IL-12p70, IL-21, CD40L, IL-10, IL-2, and IL-6 cytokine levels. Supernatant from human mKRAS- or HPV16 E7-targeted TCR-T cell, or NY-ESO-1-targeted TCR-T cell experiments were collected and stored at −80°C. The human Th17 8-plex Milliplex Map Kit (EMD Millipore, Catalog # HTH17MAG-14K) was used to quantify secreted human TNFα, IFNγ, GM-CSF, IL-12p70, IL-21, IL-10, IL-2, and IL-6 cytokine levels. Samples were run on a Luminex LX200 machine using xMAP Technology Software.

### Nanostring gene expression analysis

Lymph nodes and tumors from mice were extracted and mechanically dissociated to create a single cell suspension at indicated timepoints. Cells were counted and resuspended by vortex in 75 µL Qiagen Buffer RLT (Catalog number 79216) per 1.5×10^5^ cells. 75 µL of sample was stored in a cryovial and sent to Canopy Biosciences to perform nCounter nanostring RNA analysis utilizing Mouse Immunology panel (561 genes). Results were organized based on gene ontology pathway classification.

### Flow Cytometry

Flow cytometric analyses were performed using the 13-color CytoFLEX S (Beckman Coulter) instrument to acquire data. Data were analyzed using FlowJo (Tree star). DAPI (0.5 mg/mL, Sigma-Aldrich) or a LIVE/DEAD fixable violet fluorescent dye (Thermo Fisher Scientific) was used to exclude dead cells in all experiments. Antibodies to the murine and human proteins used for flow cytometry are listed in Table 1. All antibodies were purchased from Biolegend, Invitrogen, or BD Biosciences.

**Table 1.** Flow Cytometry Antibodies

| **Species** | **Target** | **Fluorochrome** | **Clone** | **Manufacturer** |
| --- | --- | --- | --- | --- |
| Murine | CD90.1 | Alexa Fluor 700 | OX-7 | Biolegend |
| Murine | CD3 | PE | 145-2C11 | Biolegend |
| Murine | CD4 | PerCP/Cyanine5.5 | RM4-5 | Biolegend |
| Murine | CD8 | APC | 53-6.7 | Biolegend |
| Murine | CD62L | APC/Cyanine7 | MEL-14 | Biolegend |
| Murine | CD69 | Brilliant Violet 510 | H1.2F3 | Biolegend |
| Murine | CD44 | Brilliant Violet 650 | IM7 | Biolegend |
| Murine | CD25 | PE/Cyanine7 | PC61.5 | Invitrogen/eBioscience |
| Murine | CD3 | PerCP/Cyanine5.5 | 500A2 | Biolegend |
| Murine | CD4 | Brilliant Violet 510 | RM4-5 | Biolegend |
| Murine | IFN-γ | Brilliant Violet 510 | XMG1.2 | Biolegend |
| Murine | IFN-γ | PE | XMG1.2 | Biolegend |
| Murine | TNF-α | PE/Cyanine7 | MP6-XT22 | Biolegend |
| Murine | TNF-α | FITC | MP6-XT22 | Biolegend |
| Murine | MHC II | FITC | M5/114.15.2 | Biolegend |
| Murine | CD80 | PE | 16-10A1 | Biolegend |
| Murine | CD40 | PerCP/Cyanine5.5 | 3/23 | Biolegend |
| Murine | CD11b | APC | M1/70 | Biolegend |
| Murine | CD3 | PE/Cyanine7 | 500A2 | Biolegend |
| Murine | CD11c | Brilliant Violet 510 | N418 | Biolegend |
| Murine | F4/80 | Alexa Fluor700 | BM8 | Biolegend |
| Murine | CD86 | Brilliant Violet 605 | GL-1 | Biolegend |
| Murine | CD19 | Brilliant Violet 650 | 6D5 | Biolegend |
| Murine | Granzyme B | PE/Cyanine7 | NGZB | Invitrogen/eBioscience |
| Murine | Ki67 | Brilliant Violet 605 | 16A8 | Biolegend |
| Murine | PD-1 | FITC | 29F.1A12 | Biolegend |
| Murine | CD44 | Alexa Fluor700 | IM7 | Biolegend |
| Murine | TIM-3 | Brilliant Violet 605 | RMT3-23 | Biolegend |
| Murine | LAG-3 | Brilliant Violet 650 | C9B7W | Biolegend |
| Murine | TCR beta | APC | H57-597 | Invitrogen/eBioscience |
| Human | CD3 | Alexa Fluor700 | HIT3a | Biolegend |
| Human | CD45RA | PerCP/Cyanine5.5 | HI100 | Biolegend |
| Human | CD197 (CCR7) | APC/Cyanine7 | G043H7 | Biolegend |
| Human | CD25 | Brilliant Violet 510 | BC96 | Biolegend |
| Human | CD8 | Brilliant Violet 605 | SK1 | Biolegend |
| Human | CD69 | PE/Cyanine7 | FN50 | Biolegend |
| Human | CD4 | PE | A161A1 | Biolegend |
| Human | LAG-3 | PerCP/Cyanine5.5 | 11C3C65 | Biolegend |
| Human | TIM-3 | APC/Cyanine7 | F38-2E2 | Biolegend |
| Human | PD-1 | PE/Cyanine7 | EH12.2H7 | Biolegend |

### IVIS Imaging

C57Bl/6J mice were immunized with 20 µg AMP-mKRAS-G12D or SOL-mKRAS-G12D conjugated with fluorescein carboxylic acid (FITC) or unlabeled AMP-mKRAS-G12D. Vaccines were adjuvanted with 5 nmol AMP-CpG or SOL-CpG, respectively. 24 hours post injection, inguinal and axillary lymph nodes were harvested and imaged ex vivo using the IVIS Spectrum (PerkinElmer, Waltham, MA, USA).

### ELISPOT

Spleens were collected from the murine HLA A*11:01 TCR-T cell experiment 14 days post TCR-T cell adoptive transfer. IFNγ ELISPOT was performed using the Mouse IFNγ ELISPOT^PLUS^ kit (MABTECH product # 3321-4HPW-10). ELISPOT plates were analyzed using ImmunoSpot CTL plate reader (CTL ImmunoSpot S6 Entry Analyzer) and ImmunoSpot software (ImmunoSpot Pro 7.0.23.3 Professional DC).

### Statistics

All data were plotted and all statistical analyses were performed using GraphPad Prism 8 software (La Jolla, CA). All graphs display mean values, and the error bars represent the standard deviation. No samples or animals were excluded from the analyses. Animals were randomized for the studies, and experiments were blinded to the investigator. Statistical comparisons between groups were conducted using one-way ANOVA, two-way ANOVA, or Mann-Whitney tests as indicated. Data were considered statistically significant if the P value was less than 0.05.

## Acknowledgements

**General**: The authors thank D. M. Lidgate for expert medical writing assistance, A. Bot at Capstan Therapeutics and D. J. Irvine at M.I.T. for helpful advice and discussion, and James Yang at the National Cancer Institute for provision of reagents.

## Funding

We acknowledge support from Elicio Therapeutics.

## Author contributions

D.J.D., A.M.A., and P.C.D. designed experiments, analyzed the data, and prepared the manuscript. D.J.D., A.M.A., and J.S. performed and analyzed experiments. M.P.S., A.J., and L.M.S. designed, performed, and analyzed fluorescent AMP-peptide *in vivo* experiments and prepared corresponding manuscript sections. C.M.H. reviewed the manuscript. D.J.D. and P.C.D. provided supervision and oversaw final manuscript preparation. All authors reviewed and approved the version for publication.

## Competing interests

All authors are employees of Elicio Therapeutics and, as such, receive salary and benefits, including ownership of stock and stock options from the company. D.J.D. and P.C.D. have an Amphiphile-peptide boosting of TCR-T Cells patent pending to Elicio. The authors declare no other competing interests.

## Data and materials availability

All data needed to evaluate the conclusions in the paper are present in the paper and/or the Supplementary Materials.

## Additional information

Correspondence and requests for materials should be made to P.C.D.

## Supplementary Figures

**fig. S1:**
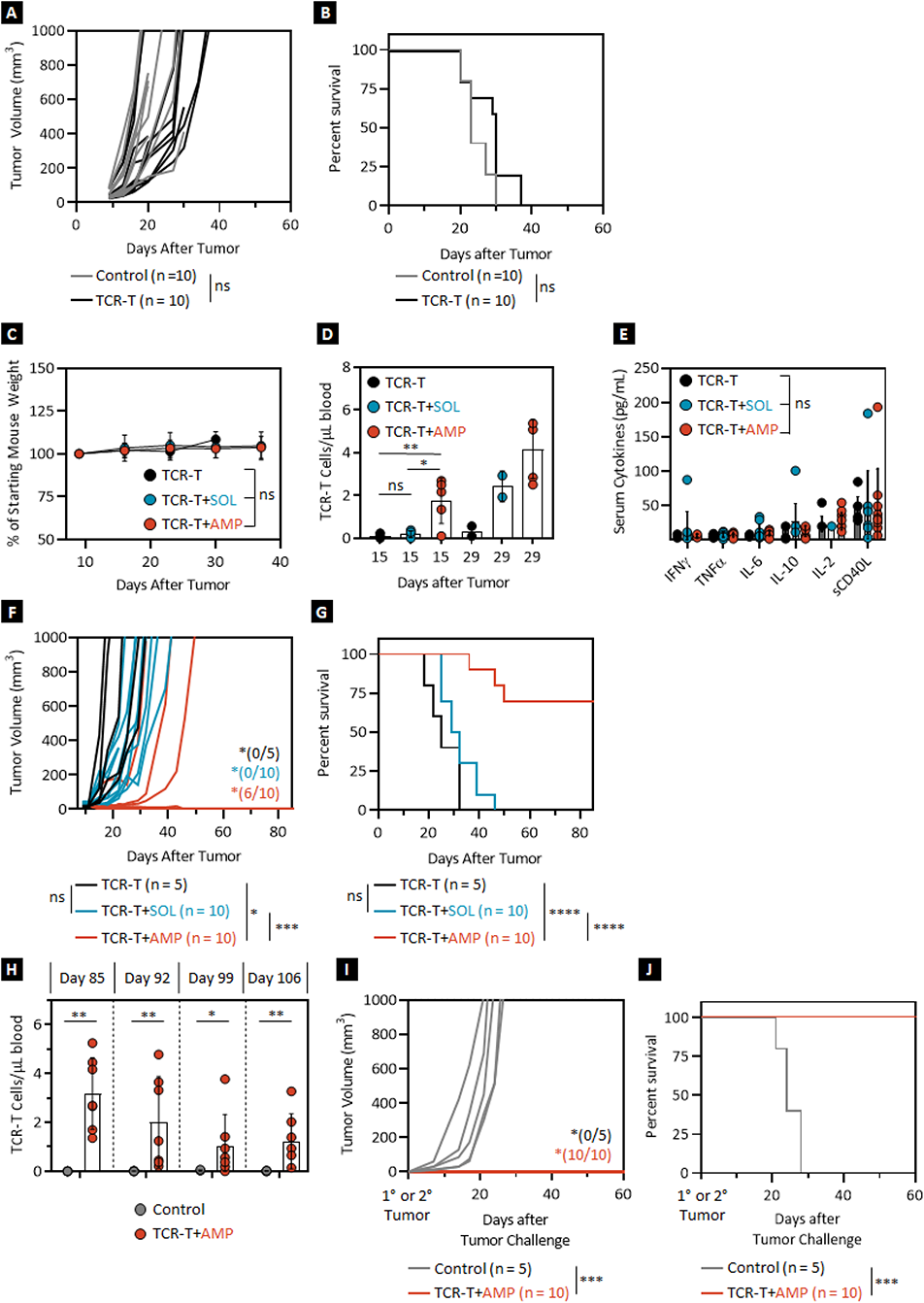
AMP-vaccination enhances TCR-T cell therapy and eradicates established solid tumors. **(A-B)** 10-day B16F10 tumor-bearing mice were irradiated (5 Gy) on day 9 with or without infusion of transduced, activated, gp100-specific TCR-T cells on day 10. **(A)** Individual tumor volume and **(B)** overall survival of treated tumor-bearing mice over time. Data shown are from 2 independent experiments. **(C-E)** 10-day B16F10 tumor-bearing mice were treated with transduced, activated, gp100-specific TCR-T cells alone or in combination with 10 µg peptide and 1 nmol adjuvant in either SOL- or AMP-gp100 vaccine regimens. **(C)** Percentage weight change of treated tumor-bearing mice during combination TCR-T cell and vaccination therapies. **(D)** Number of Thy1.1^+^ TCR-T cells per µL of peripheral blood on day 15 and 29 post tumor implantation as determined by flow cytometry. **(E)** Levels of peripheral serum cytokines on day 15 post tumor implantation as assayed by Luminex. **(F-G)** 7-day B16F10 tumor-bearing mice were treated with transduced, activated, gp100-specific TCR-T cells alone or in combination with 10 µg peptide and 1 nmol adjuvant in either SOL- or AMP-gp100 vaccine regimens. **(F)** Individual tumor volume; *indicates number of mice tumor-free at day 85 post tumor. **(G)** Overall survival over time for 7-day B16F10 tumor-bearing mice treated with transduced, activated, gp100-specific TCR-T cells alone or in combination with SOL- or AMP-gp100 vaccine regimens. Data shown are from 2 independent experiments. **(H-J)** Surviving mice that were previously implanted with 10 day-B16F10 tumors prior to treatment with transduced, activated, gp100-specific TCR-T cells in combination with 10 µg AMP-peptide and 1 nmol adjuvant or naïve control mice were rechallenged with a secondary B16F10 tumor implantation on day 85 post initial tumor challenge. **(H)** Number of Thy1.1^+^ TCR-T cells per µL of peripheral blood of mice treated with the TCR-T cells + AMP-vaccine combination in comparison to naïve control mice up to day 106 post initial tumor treatment. **(I)** Individual tumor volume; *indicates number of mice tumor-free at day 60 post tumor. **(J)** Overall survival for naïve control and surviving TCR-T cell + AMP-vaccine combination treated mice following secondary tumor challenge on day 85. Secondary tumor challenge data shown are from 2 independent experiments. TCR-T persistence data shown are means ± SDs from 2 independent experiments; p-values were determined by the Mann-Whitney Test. P-values for survival were determined by the log-rank Mantel-Cox Test, with 95% confidence interval. ****, *p* < 0.0001; ***, *p* < 0.001; **, *p* < 0.01; and *, *p* < 0.05.

**fig. S2:**
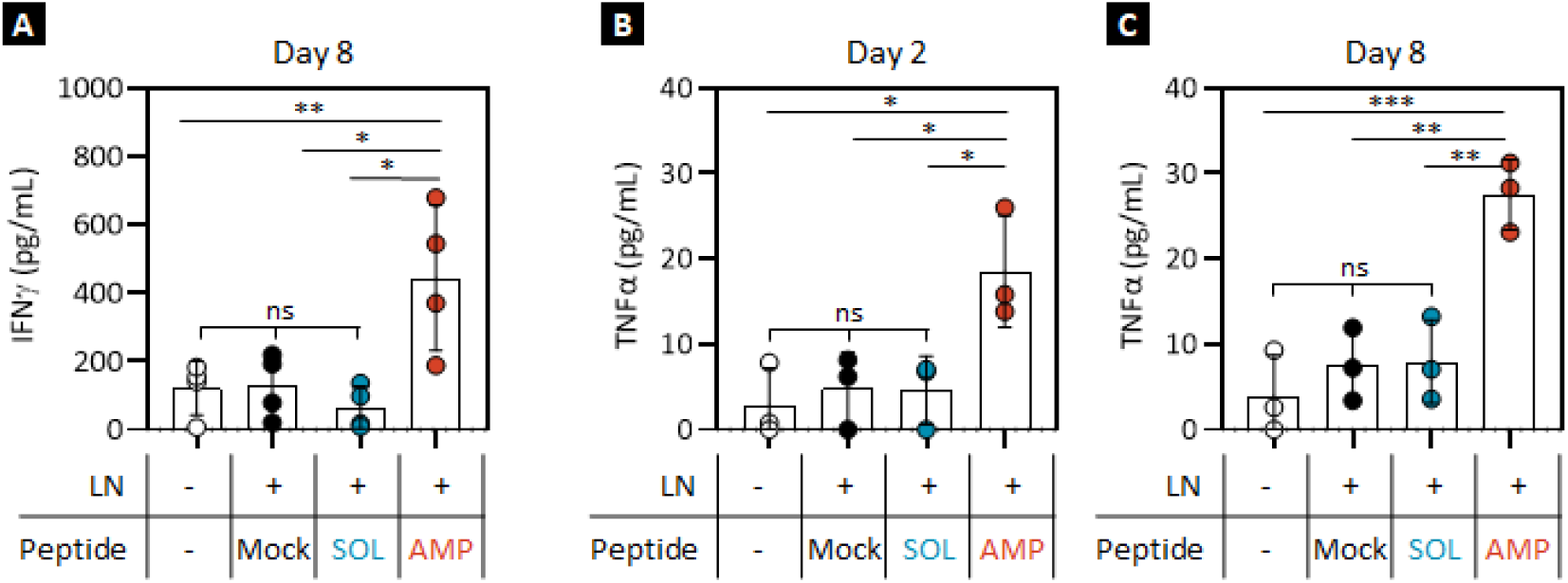
AMP-vaccines enhance TCR-T cell cytokine production ex vivo. **(A-C)** Luminex analysis of cytokines secreted by gp100-specific TCR-T cells after 24-hour co-culture with lymph node single cell suspension from mock, SOL-, or AMP-gp100-vaccinated mice. **(A)** IFNγ production of TCR-T cells 8 days following a 24-hour culture with lymph node single cell suspension from mock, SOL-, or AMP-gp100-vaccinated mice assessed by Luminex. TNFα production of TCR-T cells **(B)** 2 and **(C)** 8 days following a 24-hour culture with lymph node single cell suspensions from mock, SOL-, or AMP-gp100-vaccinated mice determined by Luminex. LN: lymph node. Data shown are means ± SDs from 3 independent experiments; p-values were determined by one-way ANOVA.***, *p* < 0.001; **, *p* < 0.01; and *, *p* < 0.05.

**fig. S3:**
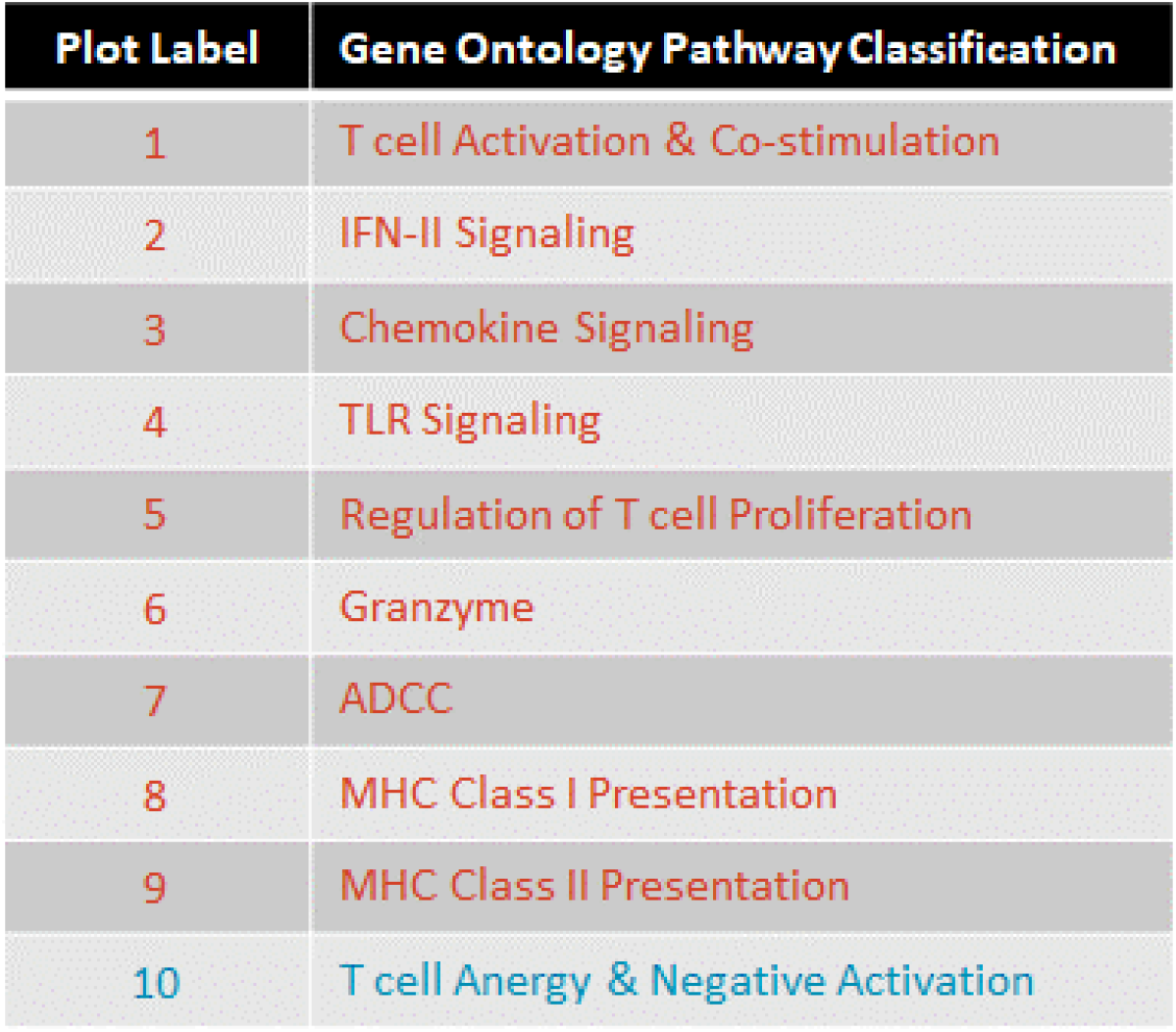
Gene ontology pathway classification for Nanostring analysis. **(A)** RNA that was extracted from lymph nodes following combination TCR-T cell therapy with mock, SOL-, or AMP-gp100-vaccination was analyzed by a 561-gene nCounter Mouse Immunology panel to perform an unbiased analysis of the lymph node transcriptome. Z-scores were calculated for the data set, and genes were clustered and plotted based upon their gene ontology classification.

**fig. S4:**
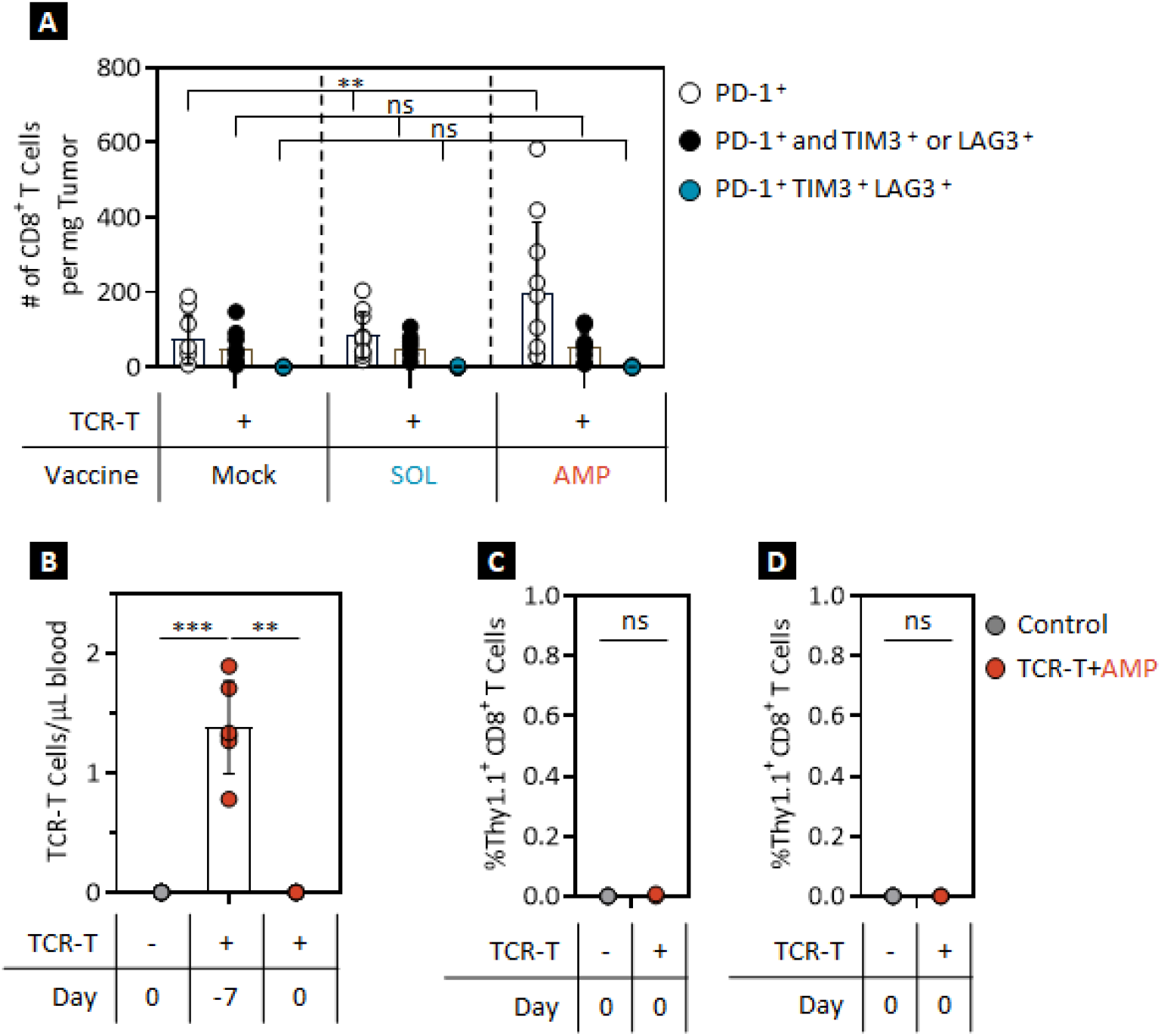
AMP-vaccination induces DC activation and inflammatory reprogramming in draining lymph nodes. **(A)** Markers of T cell exhaustion detected by flow cytometry on T cells excised from tumor samples of mock, SOL-, or AMP-gp100-vaccinated and TCR-T cell treated tumor-bearing mice. Co-expression of PD-1/TIM-3/LAG-3 exhaustion markers on tumor-infiltrating CD8^+^ T cells as determined by flow cytometry. **(B-D)** Surviving mice that were previously implanted with 10-day B16F10 tumors prior to treatment with transduced, activated, gp100-specific TCR-T cells in combination with 10 µg AMP-peptide and 1 nmol adjuvant or naïve control mice were depleted of persisting TCR-T cells by an anti-Thy1.1 antibody 7 days prior to rechallenge with a secondary B16F10 tumor implantation on day 85 post initial tumor challenge. **(B)** Number of Thy1.1^+^ TCR-T cells per µL of peripheral blood of mice before (day −7) and 1 week after (day 0) anti-Thy1.1 depletion as determined by flow cytometry. Number of Thy1.1^+^ TCR-T cells in **(C)** spleen and **(D)** lymph nodes of mice 1 week after (day 0) anti-Thy1.1 depletion as determined by flow cytometry. Data shown are means ± SDs from 2 independent experiments; p-values were determined by the Mann-Whitney test, or one- or two-way ANOVA. ***, *p* < 0.001 and **, *p* < 0.01.

**fig. S5:**
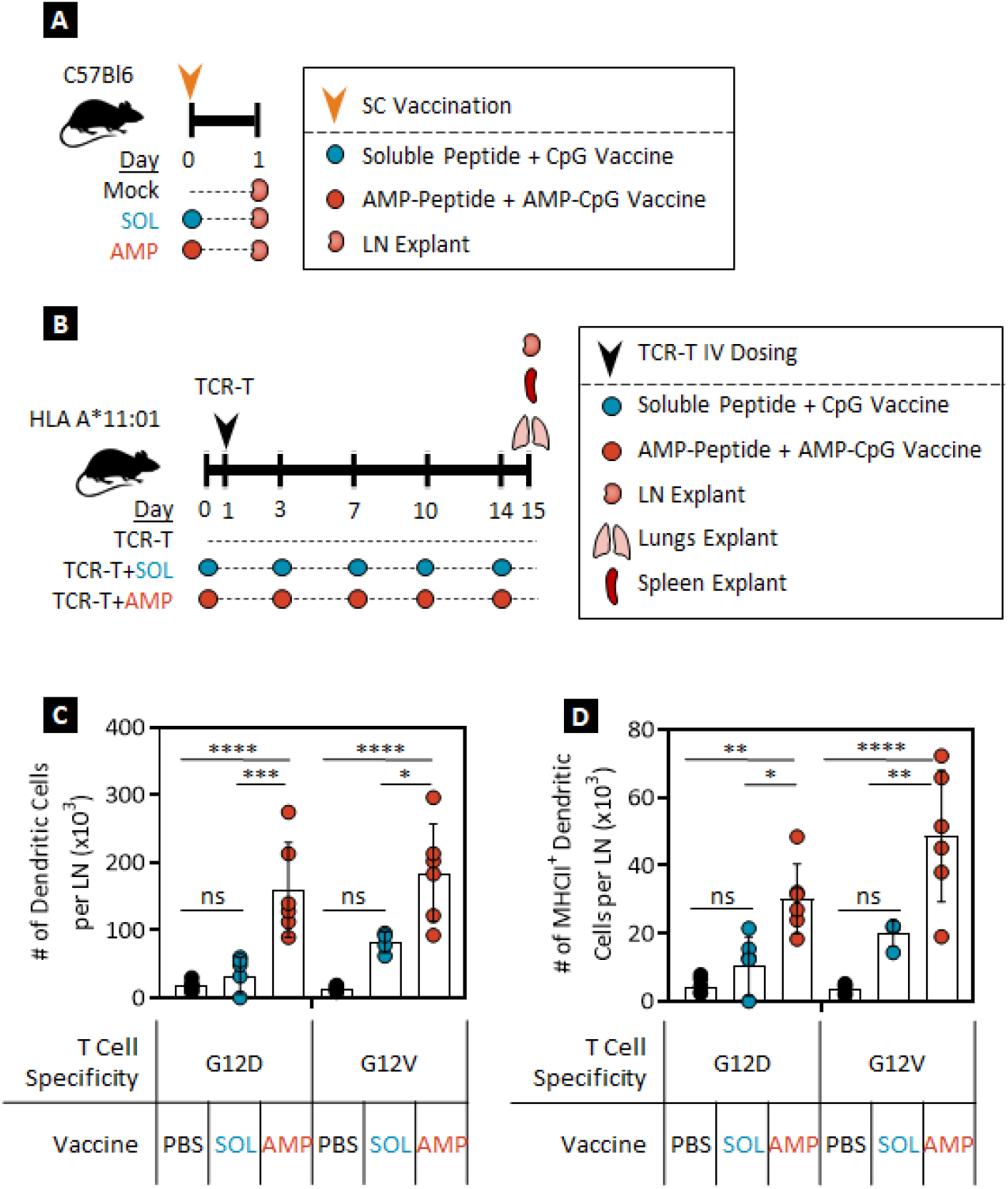
AMP-vaccination induces DC activation within lymph nodes of HLA A*11:01 transgenic mice enhancing the number of activated and functional mKRAS-specific murine TCR-T cells in vivo. **(A)** Experimental schematic depicting the immunization of C57BL/6J mice with FITC-labeled mKRAS-G12D peptides. **(B-D)** Combination therapy of HLA A*11:01-expressing mice with HLA A*11:01^+^ T cells transduced with a mKRAS-G12D-specific TCR alongside mock-, SOL-, or AMP-mKRAS-G12D peptide vaccination **(B)** Schematic of HLA A*11:01 transgenic mouse experimental protocol. **(C)** Number of CD11b/c^+^ DCs expressing **(D)** MHC Class II^+^ within inguinal lymph nodes of TCR-T cell treated and vaccinated HLA A*11:01 transgenic mice as determined by flow cytometry. LN: lymph node. Data shown are means ± SDs from 2 independent experiments; p-values were determined by the Mann-Whitney test, or one- or two-way ANOVA. ****, *p* < 0.0001; ***, *p* < 0.001; **, *p* < 0.01; and *, *p* < 0.05.

**fig. S6:**
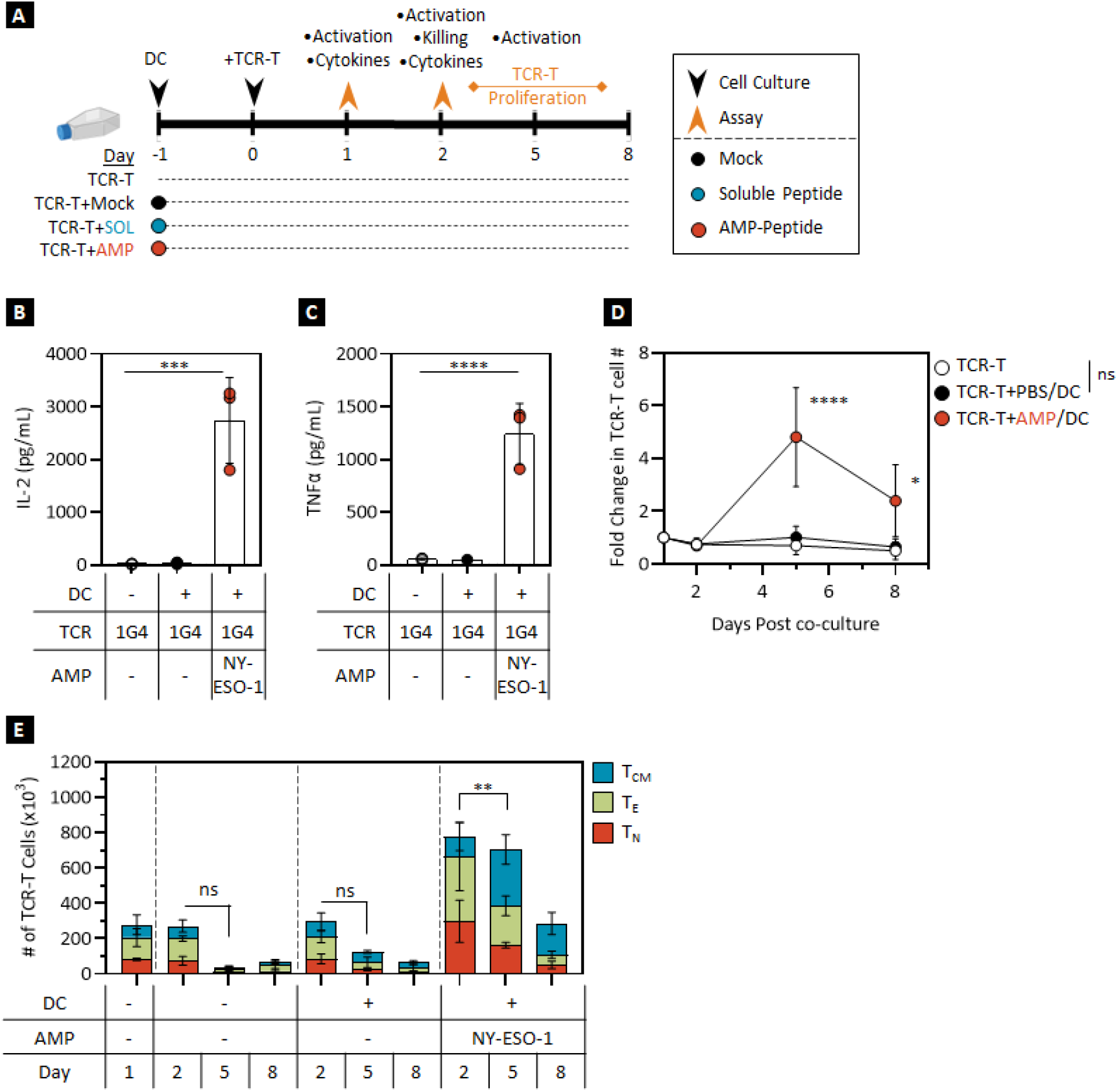
NY-ESO-1-specific human TCR-T cells are specifically activated by AMP-peptide pulsed autologous DCs in vitro. **(A-E)** NY-ESO-1-specific TCR-T cells (1G4) were cultured for 24 hours with AMP-NY-ESO-1 peptide pulsed autologous DCs prior to characterization. **(A)** Schematic of experimental protocol. TCR-T cell production of **(B)** IL-2 and **(C)** TNFα following co-culture of AMP-NY-ESO-1 peptide-pulsed autologous HLA A*02:01 human DCs with NY-ESO-1-specific HLA A*02:01 TCR-T cells as determined by Luminex. **(D**) Fold-change in TCR-T cell number following T cell:DC co-culture. **(E)** TCR-T cell phenotypic analysis following a T cell:DC co-culture as determined by flow cytometry. In vitro data shown are means ± SDs from 2 independent experiments with p-values determined by the Mann-Whitney test, one-, or two-way ANOVA. ****, *p* < 0.0001 and ***, p < 0.001.

**fig. S7:**
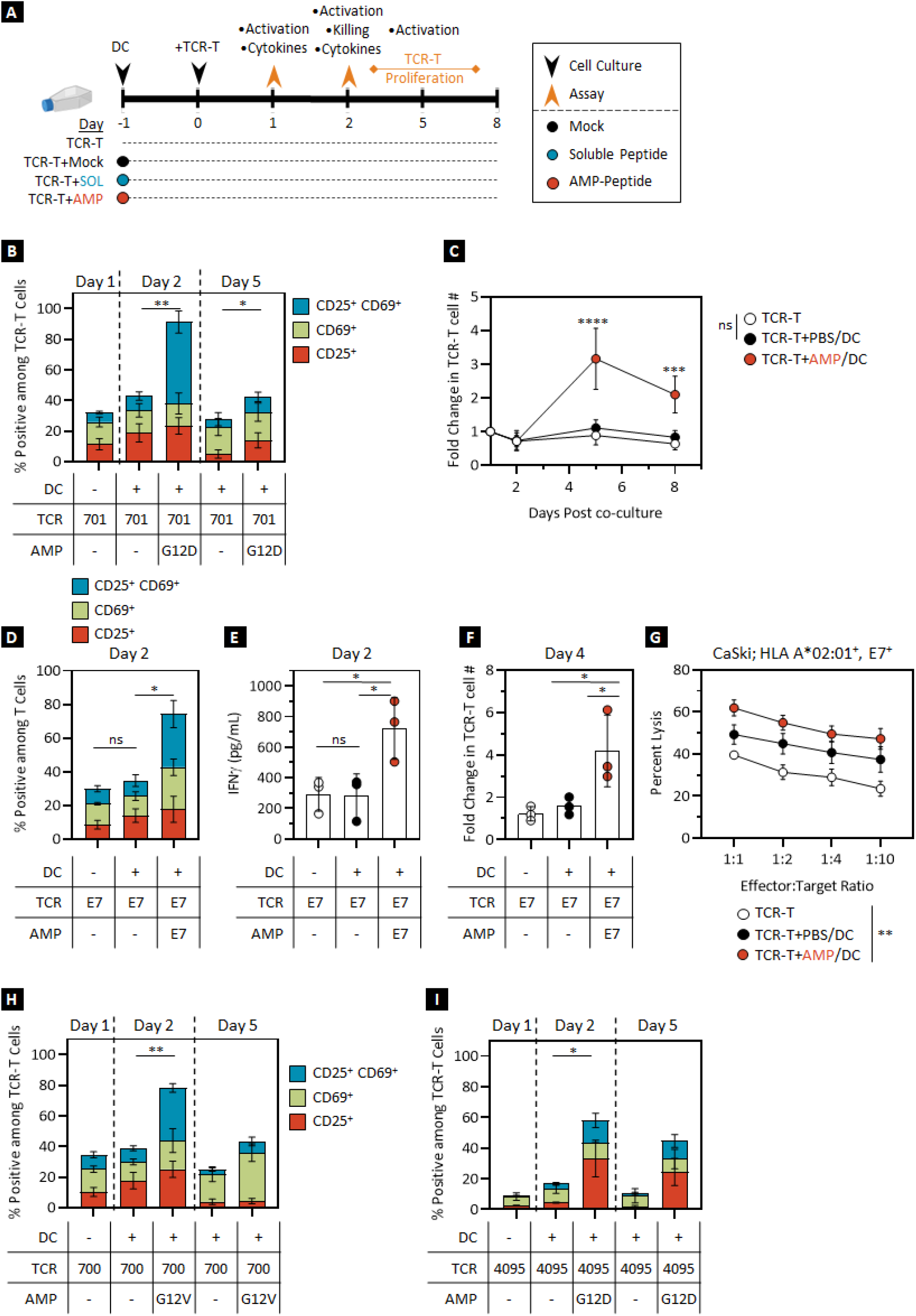
mKRAS- and HPV16 E7-specific human TCR-T cells are specifically activated by AMP-peptide-pulsed autologous DCs in vitro. **(A)** Schematic of experimental protocol. **(B-C)** mKRAS-G12D-specific HLA A*11:01 TCR-T cells (701) were cultured for 24 hours with mock or AMP-mKRAS-G12D-pulsed autologous DCs prior to characterization. TCR-T cell **(B)** CD25/CD69 expression as assessed by flow cytometry and **(C)** longitudinal fold-change in TCR-T cell number. **(D-G)** HPV16 E7-specific HLA A*02:01 TCR-T cells were cultured for 24 hours with mock or AMP-HPV16 E7-pulsed autologous DCs prior to characterization. TCR-T cell **(D)** CD25/CD69 expression as assessed by flow cytometry, **(E)** production of IFNγ by Luminex, and **(F)** fold-change in TCR-T cell number. **(G)** In vitro cytotoxicity of HPV16 E7-specific TCR-T cells with or without prior in vitro DC co-culture against HLA-matched, antigen-positive CaSki tumor cells in a luciferase killing assay at various effector-to-target ratios. **(H-I)** mKRAS-G12V-specific HLA A*11:01 (700) or mKRAS-G12D-specific HLA C*08:02 (4095) TCR-T cells were cultured for 24 hours with mock or AMP-mKRAS-G12D or -G12V-labeled autologous DCs prior to characterization. TCR-T cell CD25/CD69 expression as assessed by flow cytometry of **(H)** mKRAS-G12V-specific HLA A*11:01 TCR-T cells or **(I)** mKRAS-G12D-specific HLA C*08:02 TCR-T cells following autologous DC co-culture. In vitro data shown are means ± SDs from 2 or 3 independent experiments with p-values determined by the Mann-Whitney test, one-, or two-way ANOVA. ****, *p* < 0.0001; **, *p* < 0.01; and *, *p* < 0.05.

**fig. S8:**
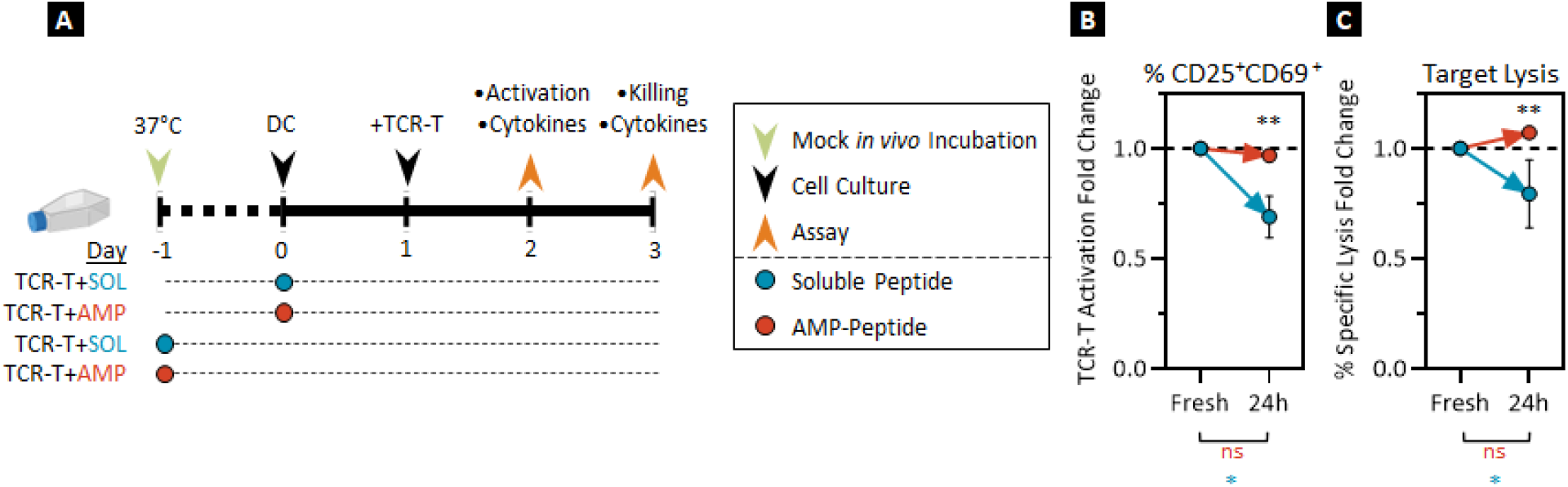
mKRAS-specific human TCR-T cells are specifically activated in vitro by autologous DCs pulsed with AMP-peptides incubated in mock in vivo conditions. **(A-C)** mKRAS-G12V-specific TCR-T cell characterization following co-culture with autologous HLA A*11:01^+^ DCs pulsed with freshly diluted SOL- and AMP-peptides or peptides that were incubated for 24 hours at 37 °C in 10% human serum to mimic in vivo conditions. **(A)** Schematic of mock in vivo experimental protocol. **(B)** TCR-T cell CD25/CD69 expression fold-change as assessed by flow cytometry. **(C)** Fold-change of in vitro cytotoxicity in a luciferase killing assay with mKRAS-specific TCR-T cells following autologous DC culture. In vitro data shown are means ± SDs from 2 or 3 independent experiments; comparisons between treatment groups shown in black, intra-group comparisons between conditions shown in corresponding colors. P-values determined by the Mann-Whitney test, one-, or two-way ANOVA.**, *p* < 0.01 and *, *p* < 0.05.

**fig. S9:**
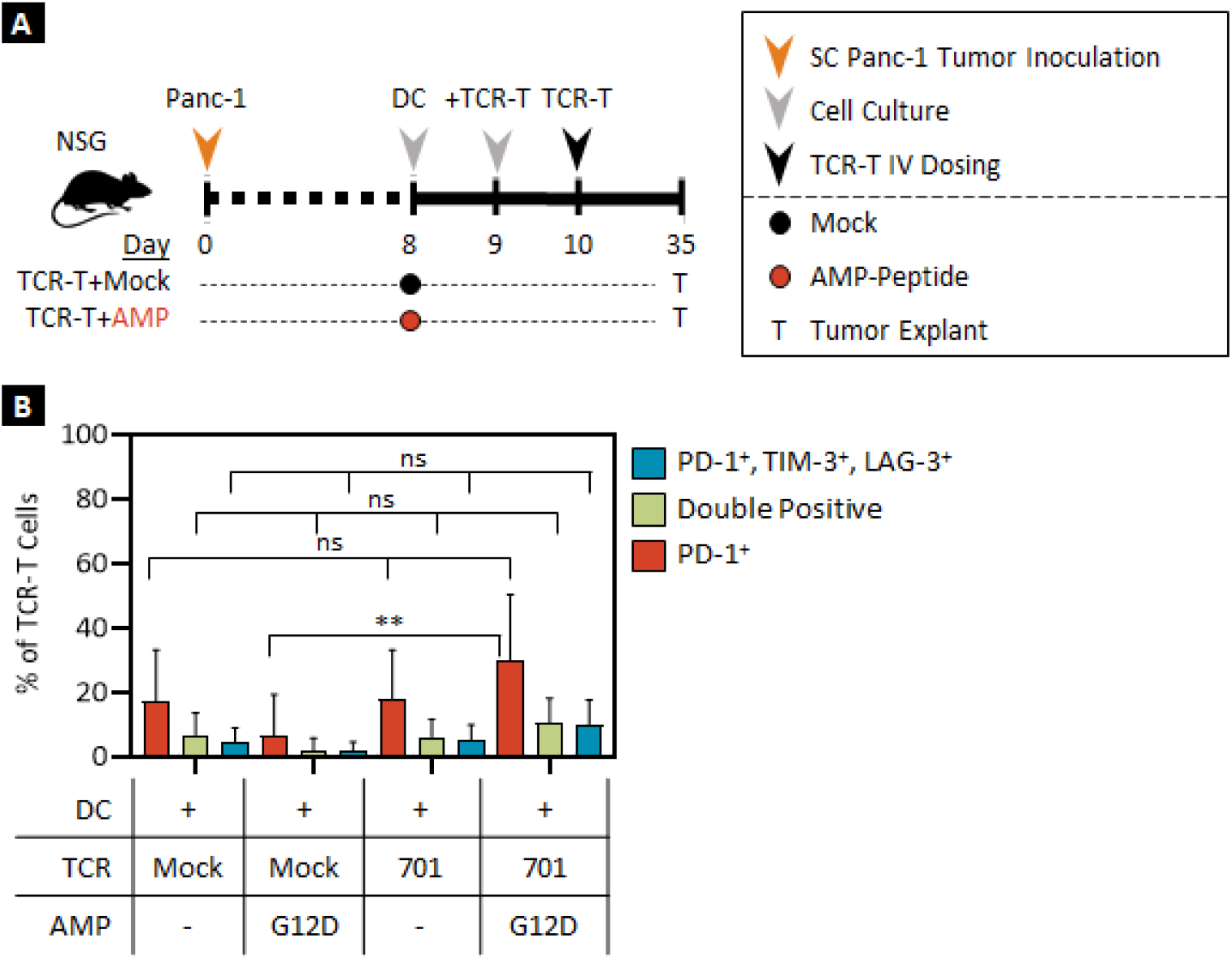
mKRAS-specific human TCR-T cells co-cultured with AMP-peptide-pulsed autologous DCs in vitro enhance in vivo anti-tumor efficacy. **(A-B)** Mock-transduced or mKRAS-G12D-specific TCR-T (701) cells were co-cultured in vitro with mock or AMP-mKRAS-G12D-pulsed DCs prior to infusion into Panc-1 tumor bearing mice to determine anti-tumor efficacy. **(A)** Schematic of experimental protocol. **(B)** Co-expression of PD-1/TIM-3/LAG-3 exhaustion markers on tumor-infiltrating mock transduced or TCR-T cells as determined by flow cytometry. In vivo data shown are means ± SDs from 1 experiment with p-values determined by two-way ANOVA. **, *p* < 0.01.

## References

1. P. F. Robbins, S. H. Kassim, T. L. N. Tran, J. S. Crystal, R. A. Morgan, S. A. Feldman, J. C. Yang, M. E. Dudley, J. R. Wunderlich, R. M. Sherry, U. S. Kammula, M. S. Hughes, N. P. Restifo, M. Raffeld, C. C. R. Lee, Y. F. Li, M. El-Gamil, S. A. Rosenberg, A pilot trial using lymphocytes genetically engineered with an NY-ESO-1-reactive T-cell receptor: Long-term follow-up and correlates with response. Clin. Cancer Res. (2015), doi:10.1158/1078-0432.CCR-14-2708.

2. S. Stevanovic, S. R. Helman, J. R. Wunderlich, M. M. Langhan, S. L. Doran, M. L. M. Kwong, R. P. T. Somerville, C. A. Klebanoff, U. S. Kammula, R. M. Sherry, J. C. Yang, S. A. Rosenberg, C. S. Hinrichs, A Phase II Study of Tumor-infiltrating Lymphocyte Therapy for Human Papillomavirus–associated Epithelial Cancers. Clin. Cancer Res. (2019), doi:10.1158/1078-0432.CCR-18-2722.

3. R. A. Morgan, M. E. Dudley, J. R. Wunderlich, M. S. Hughes, J. C. Yang, R. M. Sherry, R. E. Royal, S. L. Topalían, U. S. Kammula, N. P. Restifo, Z. Zheng, A. Nahvi, C. R. De Vries, L. J. Rogers-Freezer, S. A. Mavroukakis, S. A. Rosenberg, Cancer regression in patients after transfer of genetically engineered lymphocytes. Science (80-.). (2006), doi:10.1126/science.1129003.

4. R. A. Morgan, N. Chinnasamy, D. D. Abate-daga, A. Gros, F. Robbins, Z. Zheng, S. A. Feldman, J. C. Yang, R. M. Sherry, Q. Phan, M. S. Hughes, U. S. Kammula, A. D. Miller, C. J. Hessman, A. A. Stewart, N. P. Restifo, M. M. Quezado, M. Alimchandani, Z. Rosenberg, A. Nath, T. Wang, B. Bielekova, S. C. Wuest, A. Nirmala, F. J. Mcmahon, S. Wilde, B. Mosetter, J. Dolores, C. M. Laurencot, S. A. Rosenberg, Cancer regression and neurologic toxicity following anti-MAGE-A3 TCR gene therapy. J. Immunother. 36, 133–151 (2014).

5. M. W. Rohaan, J. H. van den Berg, P. Kvistborg, J. B. A. G. Haanen, Adoptive transfer of tumor-infiltrating lymphocytes in melanoma: a viable treatment option. J. Immunother. Cancer (2018), doi:10.1186/s40425-018-0391-1.

6. J. Zhang, L. Wang, The emerging world of TCR-T cell trials against cancer: A systematic review. Technol. Cancer Res. Treat. (2019), doi:10.1177/1533033819831068.

7. T. Isao, M. Masuya, S. Kageyama, T. Nishida, S. Terakura, M. Murata, H. Fujiwara, Y. Akatsuka, Y. Hiroaki Ikeda, Adoptive Transfer of WT1-Specific TCR Gene-Transduced Lymphocytes in Patients with Myelodysplastic Syndrome and Acute Myeloid Leukemia. Blood (2015), doi:1528-0020.

8. H. J. Stauss, S. Thomas, M. Cesco-Gaspere, D. P. Hart, S. A. Xue, A. Holler, J. King, G. Wright, M. Perro, C. Pospori, E. Morris, WT1-specific T cell receptor gene therapy: Improving TCR function in transduced T cells. Blood Cells, Mol. Dis. 40, 113–116 (2008).

9. G. Q. Phan, S. a. Rosenberg, Adoptive cell transfer for patients with metastatic melanoma: The potential and promise of cancer immunotherapy. Cancer Control. 20, 289–297 (2013).

10. S. S. Chandran, C. A. Klebanoff, T cell receptor-based cancer immunotherapy: Emerging efficacy and pathways of resistance. Immunol. Rev. (2019), doi:10.1111/imr.12772.

11. R. J. Brentjens, I. Rivière, J. H. Park, M. L. Davila, X. Wang, J. Stefanski, C. Taylor, R. Yeh, S. Bartido, O. Borquez-Ojeda, M. Olszewska, Y. Bernal, H. Pegram, M. Przybylowski, D. Hollyman, Y. Usachenko, D. Pirraglia, J. Hosey, E. Santos, E. Halton, P. Maslak, D. Scheinberg, J. Jurcic, M. Heaney, G. Heller, M. Frattini, M. Sadelain, Safety and persistence of adoptively transferred autologous CD19-targeted T cells in patients with relapsed or chemotherapy refractory B-cell leukemias. Blood. 118, 4817– 4828 (2011).

12. D. L. Porter, W. Hwang, N. V Frey, S. F. Lacey, P. A. Shaw, A. W. Loren, A. Bagg, K. T. Marcucci, A. Shen, V. Gonzalez, D. Ambrose, S. A. Grupp, A. Chew, Z. Zheng, M. C. Milone, B. L. Levine, J. J. Melenhorst, C. H. June, Chimeric antigen receptor T cells persist and induce sustained remissions in relapsed refractory chronic lymphocytic leukemia. Sci. Transl. Med. 7, 303ra139 (2015).

13. S. L. Maude, N. Frey, P. A. Shaw, R. Aplenc, D. M. Barrett, N. J. Bunin, A. Chew, V. E. Gonzalez, Z. Zheng, S. F. Lacey, Y. D. Mahnke, J. J. Melenhorst, S. R. Rheingold, A. Shen, D. T. Teachey, B. L. Levine, C. H. June, D. L. Porter, S. A. Grupp, Chimeric antigen receptor T cells for sustained remissions in leukemia. N. Engl. J. Med. (2014), doi:10.1056/NEJMoa1407222.

14. J. N. Kochenderfer, M. E. Dudley, S. H. Kassim, R. P. T. Somerville, R. O. Carpenter, S. S. Maryalice, J. C. Yang, G. Q. Phan, M. S. Hughes, R. M. Sherry, M. Raffeld, S. Feldman, L. Lu, Y. F. Li, L. T. Ngo, A. Goy, T. Feldman, D. E. Spaner, M. L. Wang, C. C. Chen, S. M. Kranick, A. Nath, D. A. N. Nathan, K. E. Morton, M. A. Toomey, S. A. Rosenberg, Chemotherapy-refractory diffuse large B-cell lymphoma and indolent B-cell malignancies can be effectively treated with autologous T cells expressing an anti-CD19 chimeric antigen receptor. J Clin Oncol. 33, 540–549 (2015).

15. X. Wang, L. L. Popplewell, J. R. Wagner, A. Naranjo, M. S. Blanchard, M. R. Mott, A. P. Norris, C. L. W. Wong, R. Z. Urak, W. C. Chang, S. K. Khaled, T. Siddiqi, L. E. Budde, J. Xu, B. Chang, N. Gidwaney, S. H. Thomas, L. J. N. Cooper, S. R. Riddell, C. E. Brown, M. C. Jensen, S. J. Forman, Phase 1 studies of central memory-derived CD19 CAR T-cell therapy following autologous HSCT in patients with B-cell NHL. Blood (2016), doi:10.1182/blood-2015-12-686725.

16. G. P. Linette, E. A. Stadtmauer, M. V. Maus, A. P. Rapoport, B. L. Levine, L. Emery, L. Litzky, A. Bagg, B. M. Carreno, P. J. Cimino, G. K. Binder-Scholl, D. P. Smethurst, A. B. Gerry, N. J. Pumphrey, A. D. Bennett, J. E. Brewer, J. Dukes, J. Harper, H. K. Tayton-Martin, B. K. Jakobsen, N. J. Hassan, M. Kalos, C. H. June, Cardiovascular toxicity and titin cross-reactivity of affinity-enhanced T cells in myeloma and melanoma. Blood (2013), doi:10.1182/blood-2013-03-490565.

17. M. de Charette, A. Marabelle, R. Houot, Turning tumour cells into antigen presenting cells: The next step to improve cancer immunotherapy? Eur. J. Cancer (2016), doi:10.1016/j.ejca.2016.09.010.

18. S. Rameshbabu, B. W. Labadie, A. Argulian, A. Patnaik, Targeting Innate Immunity in Cancer Therapy. Vaccines. 9, 1–26 (2021).

19. K. D. Moynihan, D. J. Irvine, Roles for Innate Immunity in Combination Immunotherapies. Cancer Res. 77 (2017), doi:10.1158/0008-5472.CAN-17-1340.

20. R. J. Brentjens, E. Santos, Y. Nikhamin, R. Yeh, M. Matsushita, K. La Perle, A. Quintas-Cardama, S. M. Larson, M. Sadelain, Genetically Targeted T Cells Eradicate Systemic Acute Lymphoblastic Leukemia Xenografts. Clin. Cancer Res. 13, 5426–5435 (2007).

21. R. A. Morgan, M. E. Dudley, J. R. Wunderlich, M. S. Hughes, J. C. Yang, R. M. Sherry, R. E. Royal, S. L. Topalian, U. S. Kammula, N. P. Restifo, Z. Zheng, A. Nahvi, C. R. de Vries, L. J. Rogers-Freezer, S. A. Mavroukakis, S. A. Rosenberg, Cancer regression in patients after transfer of genetically engineered lymphocytes. Science. 314, 126–129 (2006).

22. P. Johansen, D. Mohanan, J. M. Martínez-Gómez, T. M. Kündig, B. Gander, Lympho-geographical concepts in vaccine delivery. J. Control. Release. 148, 56–62 (2010).

23. H. Liu, K. D. Moynihan, Y. Zheng, G. L. Szeto, A. V. Li, B. Huang, D. S. Van Egeren, C. Park, D. J. Irvine, Structure-based programming of lymph-node targeting in molecular vaccines. Nat. 2014 5077493. 507, 519–522 (2014).

24. M. P. Steinbuck, L. M. Seenappa, A. Jakubowski, L. K. McNeil, C. M. Haqq, P. C. DeMuth, A lymph node-targeted Amphiphile vaccine induces potent cellular and humoral immunity to SARS-CoV-2. Sci. Adv. 7 (2021), doi:10.1126/SCIADV.ABE5819.

25. N. L. Trevaskis, L. M. Kaminskas, C. J. H. Porter, From sewer to saviour — targeting the lymphatic system to promote drug exposure and activity. Nat. Rev. Drug Discov. 2015 1411. 14, 781–803 (2015).

26. D. J. Irvine, M. A. Swartz, G. L. Szeto, Engineering synthetic vaccines using cues from natural immunity. Nat. Mater. 2013 1211. 12, 978–990 (2013).

27. K. D. Moynihan, C. F. Opel, G. L. Szeto, A. Tzeng, E. F. Zhu, J. M. Engreitz, R. T. Williams, K. Rakhra, M. H. Zhang, A. M. Rothschilds, S. Kumari, R. L. Kelly, B. H. Kwan, W. Abraham, K. Hu, N. K. Mehta, M. J. Kauke, H. Suh, J. R. Cochran, D. A. Lauffenburger, K. D. Wittrup, D. J. Irvine, Eradication of large established tumors in mice by combination immunotherapy that engages innate and adaptive immune responses. Nat. Med. 22, 1402 (2016).

28. K. D. Moynihan, R. L. Holden, N. K. Mehta, C. Wang, M. R. Karver, J. Dinter, S. Liang, W. Abraham, M. B. Melo, A. Q. Zhang, N. Li, S. Le Gall, B. L. Pentelute, D. J. Irvine, Enhancement of peptide vaccine immunogenicity by increasing lymphatic drainage and boosting serum stability. Cancer Immunol. Res. 6, 1025 (2018).

29. L. Ma, T. Dichwalkar, J. Y. H. Chang, B. Cossette, D. Garafola, A. Q. Zhang, M. Fichter, C. Wang, S. Liang, M. Silva, S. Kumari, N. K. Mehta, W. Abraham, N. Thai, N. Li, K. Dane Wittrup, D. J. Irvine, Enhanced CAR–T cell activity against solid tumors by vaccine boosting through the chimeric receptor. Science (80-.). 365, 162–168 (2019).

30. K. D. Moynihan, R. L. Holden, N. K. Mehta, C. Wang, M. R. Karver, J. Dinter, S. Liang, W. Abraham, M. B. Melo, A. Q. Zhang, N. Li, S. Le Gall, B. L. Pentelute, D. J. Irvine, Enhancement of peptide vaccine immunogenicity by increasing lymphatic drainage and boosting serum stability HHS Public Access. Cancer Immunol Res. 6, 1025–1038 (2018).

31. J. C. Castle, S. Kreiter, J. Diekmann, M. Löwer, N. Van De Roemer, J. De Graaf, A. Selmi, M. Diken, S. Boegel, C. Paret, M. Koslowski, A. N. Kuhn, C. M. Britten, C. Huber, Ö. Türeci, U. Sahin, Exploiting the mutanome for tumor vaccination. Cancer Res. 72, 1081–1091 (2012).

32. S. P. D’angelo, L. Melchiori, M. S. Merchant, D. Bernstein, J. Glod, R. Kaplan, S. Grupp, W. D. Tap, K. Chagin, G. K. Binder, S. Basu, D. E. Lowther, R. Wang, N. Bath, A. Tipping, G. Betts, I. Ramachandran, J. M. Navenot, H. Zhang, D. K. Wells, E. Van Winkle, G. Kari, T. Trivedi, T. Holdich, L. Pandite, R. Amado, C. L. Mackall, Antitumor activity associated with prolonged persistence of adoptively transferred NY-ESO-1c259T cells in synovial sarcoma. Cancer Discov. (2018), doi:10.1158/2159-8290.CD-17-1417.

33. D. J. Drakes, S. Rafiq, T. J. Purdon, A. V. Lopez, S. S. Chandran, C. A. Klebanoff, R. J. Brentjens, Optimization of T-cell receptor-modified T cells for cancer therapy. Cancer Immunol. Res. 18, 743–755 (2020).

34. N. F. Kuhn, T. J. Purdon, D. G. van Leeuwen, A. V. Lopez, K. J. Curran, A. F. Daniyan, R. J. Brentjens, CD40 Ligand-Modified Chimeric Antigen Receptor T Cells Enhance Antitumor Function by Eliciting an Endogenous Antitumor Response. Cancer Cell (2019), doi:10.1016/j.ccell.2019.02.006.

35. A. Kunert, M. Chmielewski, R. Wijers, C. Berrevoets, H. Abken, R. Debets, Intra-tumoral production of IL18, but not IL12, by TCR-engineered T cells is non-toxic and counteracts immune evasion of solid tumors. Oncoimmunology (2017), doi:10.1080/2162402X.2017.1378842.

36. G. Krenciute, B. L. Prinzing, Z. Yi, M. F. Wu, H. Liu, G. Dotti, I. V. Balyasnikova, S. Gottschalk, Transgenic Expression of IL15 Improves Antiglioma Activity of IL13Rα2-CAR T Cells but Results in Antigen Loss Variants. Cancer Immunol. Res. 5, 571–581 (2017).

37. L. V. Hurton, H. Singh, A. M. Najjar, K. C. Switzer, T. Mi, S. Maiti, S. Olivares, B. Rabinovich, H. Huls, M. A. Forget, V. Datar, P. Kebriaei, D. A. Lee, R. E. Champlin, L. J. N. Cooper, Tethered IL-15 augments antitumor activity and promotes a stem-cell memory subset in tumor-specific T cells. Proc. Natl. Acad. Sci. U. S. A. 113, E7788–E7797 (2016).

38. L. Tang, Y. Zheng, M. B. Melo, L. Mabardi, A. P. Castaño, Y. Q. Xie, N. Li, S. B. Kudchodkar, H. C. Wong, E. K. Jeng, M. V. Maus, D. J. Irvine, Enhancing T cell therapy through TCR-signaling-responsive nanoparticle drug delivery. Nat. Biotechnol. 36, 707– 716 (2018).

39. K. C. Conlon, E. Lugli, H. C. Welles, S. A. Rosenberg, A. T. Fojo, J. C. Morris, T. A. Fleisher, S. P. Dubois, L. P. Perera, D. M. Stewart, C. K. Goldman, B. R. Bryant, J. M. Decker, J. Chen, T. A. Worthy, W. D. Figg, C. J. Peer, M. C. Sneller, H. C. Lane, J. L. Yovandich, S. P. Creekmore, M. Roederer, T. A. Waldmann, Redistribution, hyperproliferation, activation of natural killer cells and CD8 T cells, and cytokine production during first-in-human clinical trial of recombinant human interleukin-15 in patients with cancer. J. Clin. Oncol. 33, 74–82 (2015).

40. S. K. Eskandari, I. Sulkaj, M. B. Melo, N. Li, H. Allos, J. B. Alhaddad, B. Kollar, T. J. Borges, A. S. Eskandari, M. A. Zinter, S. Cai, J. P. Assaker, J. Y. Choi, B. S. Al Dulaijan, A. Mansouri, Y. Haik, B. A. Tannous, W. J. van Son, H. G. D. Leuvenink, B. Pomahac, L. V. Riella, L. Tang, M. A. J. Seelen, D. J. Irvine, J. R. Azzi, Regulatory T Cells Engineered with TCR-Signaling-Responsive IL-2 Nanogels Suppress Alloimmunity in Sites of Antigen Encounter. Sci. Transl. Med. 12 (2020), doi:10.1126/SCITRANSLMED.AAW4744.

41. J. P. Leonard, M. L. Sherman, G. L. Fisher, L. J. Buchanan, G. Larsen, M. B. Atkins, J. A. Sosman, J. P. Dutcher, N. J. Vogelzang, J. L. Ryan, Effects of single-dose interleukin-12 exposure on interleukin-12 associated toxicity and interferon-γ production. Blood (1997).

42. H. J. Pegram, J. C. Lee, E. G. Hayman, G. H. Imperato, T. F. Tedder, M. Sadelain, R. J. Brentjens, Tumor-targeted T cells modified to secrete IL-12 eradicate systemic tumors without need for prior conditioning. Blood. 119, 4133–4141 (2012).

43. O. O. Yeku, T. J. Purdon, M. Koneru, D. Spriggs, R. J. Brentjens, Armored CAR T cells enhance antitumor efficacy and overcome the tumor microenvironment. Sci. Rep. 7, 1–14 (2017).

44. S. P. Kerkar, P. Muranski, A. Kaiser, A. Boni, L. Sanchez-Perez, Z. Yu, D. C. Palmer, R. N. Reger, Z. A. Borman, L. Zhang, R. A. Morgan, L. Gattinoni, S. A. Rosenberg, G. Trinchieri, N. P. Restifo, Tumor-specific CD8+ T cells expressing interleukin-12 eradicate established cancers in lymphodepleted hosts. Cancer Res. (2010), doi:10.1158/0008-5472.CAN-10-0735.

45. L. Zhang, S. P. Kerkar, Z. Yu, Z. Zheng, S. Yang, N. P. Restifo, S. A. Rosenberg, R. A. Morgan, Improving Adoptive T Cell Therapy by Targeting and Controlling IL-12 Expression to the Tumor Environment. Mol. Ther. 19, 751–759 (2011).

46. H. Mizuguchi, IRES-Dependent Second Gene Expression Is Significantly Lower Than Cap-Dependent First Gene Expression in a Bicistronic Vector. Mol. Ther. 1, 376–382 (2000).

47. X. Wang, C. W. Wong, R. Urak, A. Mardiros, L. E. Budde, W. C. Chang, S. H. Thomas, C. E. Brown, C. La Rosa, D. J. Diamond, M. C. Jensen, R. Nakamura, J. A. Zaia, S. J. Forman, CMVpp65 vaccine enhances the antitumor efficacy of adoptively transferred CD19-redirected CMV-specific T cells. Clin. Cancer Res. 21, 2993 (2015).

48. M. Tanaka, H. Tashiro, B. Omer, N. Lapteva, J. Ando, M. Ngo, B. Mehta, G. Dotti, P. R. Kinchington, A. M. Leen, C. Rossig, C. M. Rooney, Vaccination targeting native receptors to enhance the function and proliferation of chimeric antigen receptor (CAR)-modified T cells HHS Public Access. Clin Cancer Res. 23, 3499–3509 (2017).

49. C. Y. Slaney, B. Von Scheidt, A. J. Davenport, P. A. Beavis, J. A. Westwood, S. Mardiana, D. C. Tscharke, S. Ellis, H. M. Prince, J. A. Trapani, R. W. Johnstone, M. J. Smyth, M. W. Teng, A. Ali, Z. Yu, S. A. Rosenberg, N. P. Restifo, P. Neeson, P. K. Darcy, M. H. Kershaw, Dual-specific chimeric antigen receptor T cells and an indirect vaccine eradicate a variety of large solid tumors in an immunocompetent, self-antigen setting. Clin. Cancer Res. 23, 2478–2490 (2017).

50. J. Gust, K. A. Hay, L. A. Hanafi, D. Li, D. Myerson, L. F. Gonzalez-Cuyar, C. Yeung, W. C. Liles, M. Wurfel, J. A. Lopez, J. Chen, D. Chung, S. H. Baker, T. Ozpolat, K. R. Fink, S. R. Riddell, D. G. Maloney, C. J. Turtle, Endothelial activation and blood–brain barrier disruption in neurotoxicity after adoptive immunotherapy with CD19 CAR-T cells. Cancer Discov. 7, 1404–1419 (2017).

51. J. N. Kochenderfer, M. E. Dudley, S. A. Feldman, W. H. Wilson, D. E. Spaner, I. Maric, M. Stetler-Stevenson, G. Q. Phan, M. S. Hughes, R. M. Sherry, J. C. Yang, U. S. Kammula, L. Devillier, R. Carpenter, D. A. N. Nathan, R. A. Morgan, C. Laurencot, S. A. Rosenberg, B-cell depletion and remissions of malignancy along with cytokine-associated toxicity in a clinical trial of anti-CD19 chimeric-antigen-receptor–transduced T cells. Blood. 119, 2709 (2012).

52. J. L. Gulley, R. A. Madan, R. Pachynski, P. Mulders, N. A. Sheikh, J. Trager, C. G. Drake, Role of Antigen Spread and Distinctive Characteristics of Immunotherapy in Cancer Treatment (2017), doi:10.1093/jnci/djw261.

53. X. Wang, C. W. Wong, R. Urak, A. Mardiros, L. E. Budde, W.-C. Chang, S. H. Thomas, C. E. Brown, C. La Rosa, D. J. Diamond, M. C. Jensen, R. Nakamura, J. A. Zaia, S. J. Forman, S. J. Forman, X. Wang, C. W. Wong, R. Urak, A. Mardiros, L. E. Budde, S. H. Thomas, C. E. Brown, C. La Rosa, D. J. Diamond, M. C. Jensen, R. Nakamura, J. A. Zaia, CMVpp65 vaccine enhances the antitumor efficacy of adoptively transferred CD19-redirected CMV-specific T cells HHS Public Access. Clin Cancer Res. 21, 2993–3002 (2015).

54. K. Reinhard, B. Rengstl, P. Oehm, K. Michel, A. Billmeier, N. Hayduk, O. Klein, K. Kuna, Y. Ouchan, S. Wöll, E. Christ, D. Weber, M. Suchan, T. Bukur, M. Birtel, V. Jahndel, K. Mroz, K. Hobohm, L. Kranz, M. Diken, K. Kühlcke, Ö. Türeci, U. Sahin, An RNA vaccine drives expansion and efficacy of claudin-CAR-T cells against solid tumors. Science (80-.). 367, 446–453 (2020).

55. B. Y. Jin, T. E. Campbell, L. M. Draper, S. Stevanović, B. Weissbrich, Z. Yu, N. P. Restifo, S. A. Rosenberg, C. L. Trimble, C. S. Hinrichs, Engineered T cells targeting E7 mediate regression of human papillomavirus cancers in a murine model. JCI insight (2018), doi:10.1172/jci.insight.99488.

